# Adenoviral chromatin organization primes for early gene activation

**DOI:** 10.1101/2020.07.07.190926

**Authors:** Uwe Schwartz, Tetsuro Komatsu, Claudia Huber, Floriane Lagadec, Conradin Baumgartl, Elisabeth Silberhorn, Margit Nuetzel, Fabienne Rayne, Eugenia Basyuk, Edouard Bertrand, Michael Rehli, Harald Wodrich, Gernot Laengst

## Abstract

Within the virion, adenovirus DNA associates with the virus-encoded, protamine-like structural protein pVII. Whether this association is organized, and how genome packaging changes during infection and subsequent transcriptional activation is currently unknown. Here, we combined RNA-seq, MNase-seq, ChIP-seq and single genome imaging during early adenovirus infection to unveil the structure- and time-resolved dynamics of viral chromatin changes as well as their correlation with gene transcription. Our MNase mapping data indicates that the viral genome is arranged in precisely positioned nucleoprotein particles (Adenosomes), like nucleosomes. We identified 238 Adenosomes, being positioned by a DNA sequence code and protecting about 60 to 70bp of DNA. The incoming genome is more accessible at early gene loci that undergo additional chromatin de-condensation upon infection. Histone H3.3 containing nucleosomes specifically replace pVII at distinct genomic sites and at the transcription start sites of early genes. Acetylation of H3.3 is predominant at the transcription start sites, preceding transcriptional activation. Based on our results we propose a central role for the viral pVII nucleoprotein-architecture, which is required for the dynamic structural changes during early infection, including the regulation of nucleosome assembly prior to transcription initiation. Our study thus may aid the rational development of recombinant adenoviral vectors exhibiting sustained expression in gene therapy.

## INTRODUCTION

Most DNA viruses translocate their genomes into the nucleus of infected target cells to initiate viral gene expression and genome replication. During cytoplasmic trafficking, genomes are highly compacted inside the viral capsid to protect them from nucleic acid sensing molecules. It is generally assumed that genomes are decondensed upon entering the nucleus and thereby activated for viral gene expression and replication. Replicated genomes are then re-condensed for packaging into progeny. Alternatively, nuclear viral genomes could undergo different parallel fates, i.e. one part serving as transcription unit, whereas others are replicated and/or packaged (Brison et al., 1979). While these remarkable and reversible genome functions are highly specialized and efficient, little is known about the structural organization of the genome, its dynamic changes or the regulatory cues initiating structural transitions during virus entry.

Adenoviruses (Ad) are prototypic DNA viruses containing a linear double stranded DNA genome of 30-40 kb that is incorporated into a ∼ 90 nm non-enveloped icosahedral capsid shell. Ad enter cells by receptor-mediated endocytosis. Stepwise entry cues liberate the internal capsid protein VI and permit the virus to escape the endosomal compartment (Greber et al., 1993; Wickham et al., 1993; Wiethoff et al., 2005). Cytoplasmic capsids are transported to the nuclear pore complex (NPC) where they dock, disassemble and release the viral genome for nuclear import (Cassany et al., 2015; Strunze et al., 2011). Within hours after cell entry, nuclear genomes initiate gene expression from the immediate early promoter. In a simplified view this promoter encodes the E1A gene, encoding a *trans*-activator driving early genes responsible for cell and immune modulation (E1B, E3, E4) and replication (E2). In contrast activation of late genes (L1-L5), encoding most structural proteins, requires prior genome replication (Akusjarvi, 2008; Flint, 1982).

Packaged Ad genomes are devoid of histones. Instead they associate with basic core proteins VII (pVII), V (pV) and polypeptide µ (Brown et al., 1975; Corden et al., 1976). Protein VII is the most tightly bound and most abundant protein with over 500 copies per virion and responsible for packaging the viral genome into a structure appearing as irregular “beads-on-a-string” by electron microscopy (Benevento et al., 2014). Protein VII exhibits sequence similarities to protamines and may use its positive charges to neutralise and compact the viral DNA (Anderson et al., 1989; Johnson et al., 2004; Keller et al., 2002). Cellular factors and/or post-transcriptional modification control the association of pVII with viral genomes in the producer cell prior or during genome packaging. A recent study showed that viral genomes can be packaged without pVII, but that its presence is a prerequisite for infection (Avgousti et al., 2016; Genoveso et al., 2020; Inturi et al., 2017; Mun and Punga, 2018; Ostapchuk et al., 2017; Samad et al., 2012). The genomic binding sites of pVII as well as the structural organisation it imposes on viral DNA, inside the virion, post infection and post nuclear genome delivery, remain elusive (Giberson et al., 2011; Mirza and Weber, 1982). Protein VII stays associated with the viral genome imported to the nucleus, while pV dissociates from the viral DNA following ubiquitylation (Puntener et al., 2011). The fate of the µ-peptide is not known.

Cellular TAF-Iβ/SET associates with pVII on nuclear viral genomes and is thought to stimulate transcription and prevent the activation of DNA damage responses (Avgousti et al., 2017; Giberson et al., 2011; Haruki et al., 2003; Komatsu et al., 2015). Protein VII was also shown to functionally interact with the viral *trans*-activator E1A, pinpointing to a direct role in transcriptional activation (Johnson et al., 2004). Some studies suggested that pVII remains attached throughout the early phase of infection while others argue for a gradual and at least partial removal from the incoming genome (Haruki et al., 2003; Johnson et al., 2004; Komatsu et al., 2011, 2015; Xue et al., 2005). Conflicting reports postulated that pVII turnover either requires transcription (Chen et al., 2007) or occurs independent of transcription, at least on viral vector genomes (Ross et al., 2011). In contrast chromatin immune precipitation (ChIP) analysis proposed that pVII remains associated with viral chromatin for several hours (Chatterjee et al., 1986; Haruki et al., 2003; Johnson et al., 2004; Komatsu et al., 2011; Xue et al., 2005). In addition, cellular histones may associate with incoming genomes prior to replication (Giberson et al., 2018; Ross et al., 2011).

Functional chromatin extraction using high and low concentrations of micrococcal nuclease (MNase) to hydrolyze cellular chromatin, combined with high-throughput sequencing, is a way to analyse protein binding to DNA and to reveal dynamic changes in chromatin structure (Mueller et al., 2017; Schwartz et al., 2018). In this study we combined time-resolved MNase, ChIP and RNA sequencing as well as single molecule imaging to correlate chromatin structure changes with transcription at very early time points of adenoviral infection. This approach revealed the specific nucleoprotein architecture of the virus, which dynamically changes during early infection. Protein VII positioning is driven by sequence and is specifically organized at early genes, correlating with the early de-condensation of these regions upon infection. Defined pVII nucleoprotein complexes are replaced by nucleosomes containing H3.3 on early gene promoters and few additional genomic sites. pVII remodeling, nucleosome assembly and the specific acetylation of the promoter nucleosomes are preceding transcriptional activation. The time resolved single molecule imaging of transcribing genomes support our findings, identifying the morphological correlate at the single genome level at subcellular resolution. The defined positioning of pVII on the viral genome and its functional involvement in dynamic chromatin rearrangement during infection, suggest a key role in regulating the viral life cycle. We identify a DNA sequence driven adenovirus nucleoprotein architecture, which may aid the design of improved Ad vectors for gene-therapy or vaccination.

## RESULTS

### Time resolved analysis of the early adenovirus transcriptome by RNA-sequencing

To investigate the early spatio-temporal reorganization of viral chromatin and transcription activation we infected H1299, a p53-deficient non-small cell lung carcinoma cell line, or U2OS, a human osteosarcoma cell line, with a partially E3-deleted but replication competent human adenovirus C5 (HAd-C5). The inoculum was replaced with fresh medium after 30 min, defining time point zero of the experimental setup (Fig. 1A). On time point 0, 0.5, 1, 2, and 4 hours post infection (hpi) infected cells were subjected to immune fluorescence analysis (IF), RNA-sequencing (RNA-seq), ChIP-seq and functional chromatin extraction combined with high-throughput sequencing of protected DNA fragments (MNase-seq) (Fig. 1A). To verify the progression of infection, cells were first analysed by immunofluorescence (Fig. S1A) for the internal capsid protein VI (pVI), which can be transiently detected during endosomal escape (Martinez et al., 2015), while nuclear import is marked by accessibility of the genome bound pVII in the nucleus (Komatsu et al., 2015). Protein VI and pVII quantification showed that endosomal escape occurred within the first 30 min (Fig. S1A and S1B) followed by rapid nuclear delivery of viral genomes starting at 0.5 hpi, with most genomes being imported 1 hpi (Fig. S1A and S1C). The data is confirming a synchronized infection covering pre- and post-nuclear genome delivery phases.

**Figure 1.**
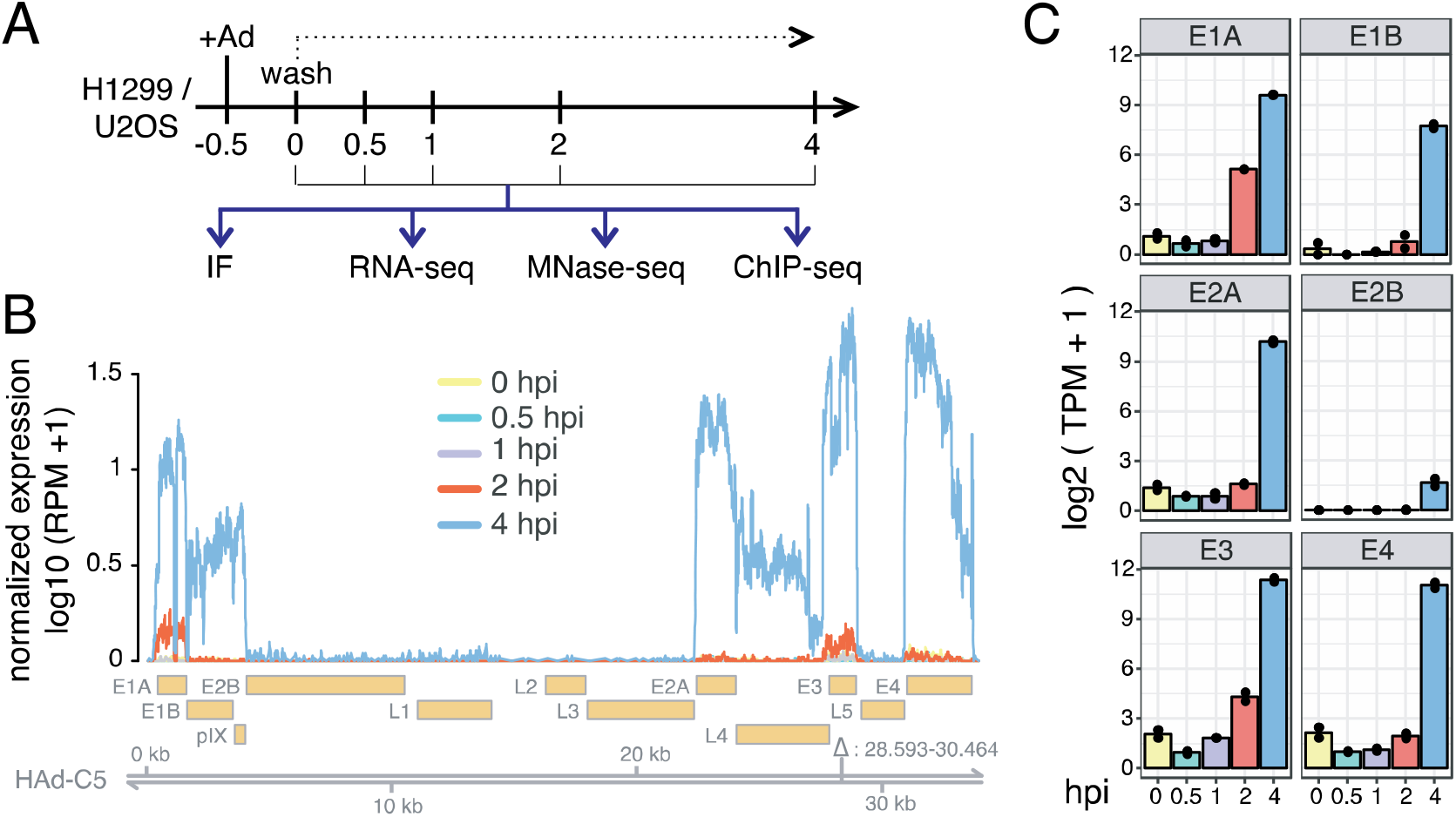
Spatio-temporal analysis of adenovirus transcription using RNA-seq. **A)** Experimental design. Virus was added (+Ad) and removed (Wash) after 30 min and cell were processed as indicated (hpi = hours post infection) using immunofluorescence (IF), RNA sequencing (RNA-seq), nucleosome sequencing (MNase-seq) and ChIP sequencing (ChIP-seq). **B)** Profile illustrating Ad genome coverage by RNA-seq reads. Reads were normalized to RPM (reads per million mapped reads) and the average of two replicates is shown. Genomic location and underlying gene annotation of HAd-C5 (NCBI accession: AY339865.1) are indicated at the bottom. The area deleted in the HAd-C5 strain is indicated by the triangle. **C)** Transcript abundance estimation of early Ad genes. TPMs (Transcripts Per Kilobase Million) for early expressed genes were calculated at each timepoint. The barplots show the average of two replicates and each replicate is indicated by a black dot.

To investigate viral transcription, we performed mRNA sequencing of two independent infection time-courses yielding on average 26 million reads per sample (Table S1). The reads were mapped to an extended version of the human genome containing the HAd-C5 reference genome and gene annotation. The first mature Ad transcripts were detected at 2 hpi, and strongly increased levels were observed 4 hpi (Fig. 1B and 1C; whole genome profile shown as individual replicates in Fig. S2 and quantification in Table S1). Consistent with the spatial organisation of the Ad transcription program (Crisostomo et al., 2019; Glenn and Ricciardi, 1988), transcription is initiated at both ends of the Ad genome covering the early genes (Fig. 1B). We detected E1A as the first viral transcript (Nevins et al., 1979), along with significant levels of E3 at 2 hpi (Fig. 1B and 1C). At 4 hpi viral transcription is markedly enhanced with high levels of transcripts originating from all early genes and significant signals arising from the E2A region. Taken together we observed an efficient utilization of the host cell transcription machinery within the first hours of infection, revealing a transcriptional competent viral template.

### *In situ* detection of E1A mRNA identifies transcribing Ad genomes

To directly correlate the progression of infection and the onset of transcription, the import of genome associated pVII and the expression of the E1A gene was monitored simultaneously by immunofluorescence (Fig. 2A). We used RNAscope technology for the direct detection of accumulating individual viral transcripts at the single cell level (Pheasant et al., 2018). A specific set of probes for the E1A mRNA was designed (ACDBio) and validated on cells infected with either HAd-C5 virus or an E1-deleted vector (Ad5-GFP). At 4 hpi specific E1A transcripts could be readily detected in virus-but not in E1-deleted GFP expressing viral vector infected cells (Fig. 2B). Protein VII immunofluorescence analysis showed that most genomes were imported before 1 hpi with no significant additional import after this time point (Fig. 2C). While the nuclear import of genomes was saturated 1 hpi, RNA-seq and RNAscope revealed that E1A transcripts were detected only at 2 hpi and accumulated at 4 hpi (Fig. 2D and Fig. 1C). Ad transcription initiation and mRNA synthesis exhibited a significant lag time after nuclear entry, suggesting a requirement for remodelling processes prior to the assembly of the transcription machinery.

**Figure 2.**
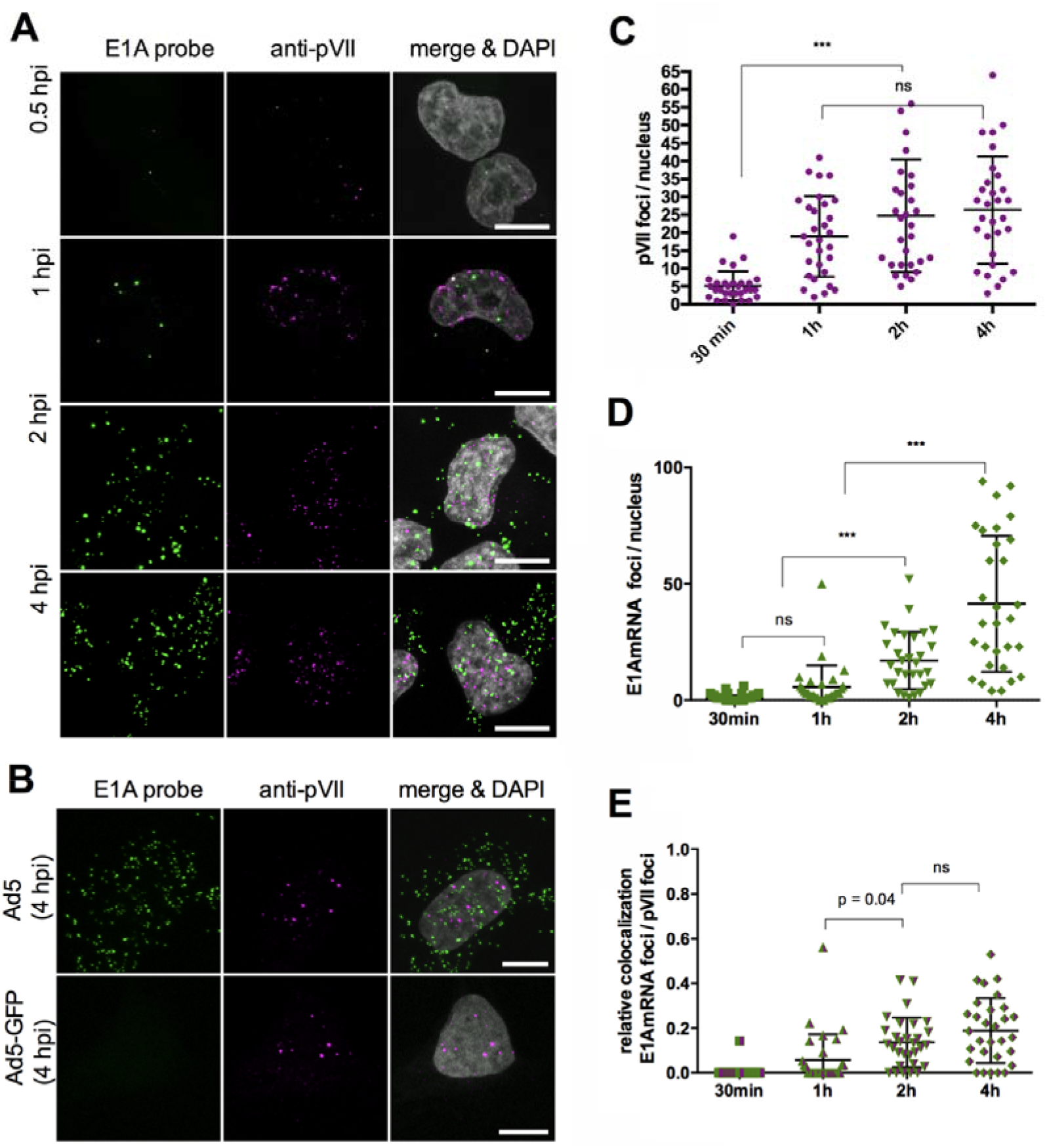
E1A transcription at the single cell level. **A)** Representative confocal images of cells infected with HAd-C5 at different time points as indicated. Detection of E1A transcript with specific RNAscope probes (left column, green signal), genomes using anti-pVII antibodies (middle column, magenta signal) and merged with DAPI detecting the nucleus (right column, grey signal). Scale bar is 10µm. **B)** Representative confocal images of cells infected with HAd-C5 (Ad5, top row) or an E1-deleted HAd-C5 vector (Ad5-GFP, bottom row). Scale bar is 10µm. **C)** Quantification of nuclear pVII foci over time as indicated on the x-axis. The distribution of normalized pVII foci at each time point is shown as scatter plot (Mean +/-SD, n = 30 cells per time point). **D)** Graph as in (C) showing the distribution of nuclear E1A transcript foci per cell as scatter plot (Mean +/-SD, n = 30 cells per time point). **E)** Scatter plot showing the distribution of nuclear pVII foci (C) coinciding with E1A transcript foci (D) (Mean +/-SD, n = 30 nuclei per time point). The number of E1A positive genomes was normalized by the total number of genomes per nucleus. Statistical analysis for C-E via one-way ANOVA multi-comparison test.

E1A transcript accumulation could be related to the continued expression from a stable pool of transcriptionally active genomes or by an increasing number of genomes initiating transcription over time. We quantified the colocalization of nuclear genomes with nuclear E1A transcripts to discriminate between both possibilities. Colocalizations inside the nucleus resulting from association by chance are expected to increase over time when active genome numbers remain constant and E1A nuclear transcript numbers increase over time. However, our results showed that E1A positive nuclear genomes rapidly plateau at ∼20% despite the strong accumulation of nuclear E1A transcripts (Fig. 2E). It is shown that only a relatively small subpopulation of imported genomes contributes to early viral gene expression until 4 hpi, which was also observed in a recent study (Suomalainen et al., 2020). These observed ∼20% transcribing viral genomes could also be the result of a larger proportion of genomes that undergo transcriptional bursting. This interpretation would not contradict the observed results.

### *In vivo* detection of E1A mRNA identifies a subpopulation of transcribing Ad genomes

The RNAscope analysis confirmed RNA-seq results and indicated, that active viral genomes undergo a lag phase in the nucleus, probably acquiring structural changes that allow to activate early gene expression. To confirm this observation and further characterize transcribing and non-transcribing genomes we developed an imaging system for living cells. To mark E1A transcripts in living cells we inserted 24x MS2 binding sites into the 3’UTR of the E1A gene of a replication competent virus. MS2 binding sites are short stem-loop structures derived from the MS2 bacteriophage and can be detected by binding of multiple copies of fluorophore tagged MS2 binding protein (i.e. MS2BP-NLS-GFP) forming a fluorescent spot *in situ* (Bertrand et al., 1998). Production and amplification of the virus resulted in normal yields with unaltered infectivity or protein composition compared to the parental virus (data not shown). To detect Ad genomes in living cells we used our previously established live cell imaging system based on fluorescent TAF-Iβ, which associates with the pVII of incoming genomes, thereby generating a fluorescent spot *in situ* (Komatsu et al., 2015). A stable mCherry-TAF-Iβ expressing cell line was transduced with an expression vector for MS2BP-NLS-GFP to generate double fluorescent stable cell lines. Cells were left either uninfected, or infected with untagged replicative control virus, or infected with E1A-MS2 tagged replicative virus and at ∼ 4 hpi analysed by live cell imaging at high temporal resolution (1 frame/sec) (Movies S1-S4). To display the results in a single image we superimposed individual movie frames (60 sec t-projections, Fig. 3A). In non-infected control cells both, mCherry-TAF-Iβ and MS2BP-NLS-GFP localized to the nucleoplasm, with some unspecific enrichment of the MS2 signal in nucleoli (Fig. 3A top row, taken from Movie S1). When cells were infected with non-MS2 containing control virus, genomes formed mCherry-TAF-Iβ fluorescent spots in the nucleoplasm with limited confined motion as previously described (Komatsu et al., 2015), while the MS2 signal remained unchanged compared to non-infected cells (Fig. 3A 2^nd^ row taken from Movie S2). In contrast cells infected with virus expressing MS2-repeat-tagged E1A mRNA formed mCherry-TAF-Iβ spots indicating viral genomes but also rapidly moving GFP spots visible inside the nucleoplasm and in the cytoplasm (Movie S3/S4). GFP spots accumulated in most cells around 3-5 hpi in agreement with the RNAscope experiment. Upon merging the mCherry-TAF-Iβ channel (viral genomes) with the MS2BP-NLS-GFP channel (viral E1AmRNA) using t-projections, some genomes were clearly double positive suggesting transcribing genomes (Fig. 3A 3^rd^ and 4^th^ row, taken from Movie S3 and S4). Similar observations were made when we used a different MS2BP construct (MS2BP-NES-GFP) predominantly localizing to the cytosol or when we lowered the input MOI resulting in fewer (<10) genome copies per imaging field (Fig. S3). To quantify the number of E1A mRNA positive genomes we performed kymograph analysis using dual-channel t-projections of individual cells. This approach was possible because the genome position was stable over time allowing to draw a line plot through all nuclear genomes using the signal of mCherry-TAF-Iβ. When plotted as kymograph over time each genome is represented as a red line. Genomes positive for the MS2BP-NLS-GFP are represented as a green line and scored as transcribing genomes (Fig. 3B). Kymograph analysis of several cells showed that ∼20% of detected genomes inside a given nucleus are positive for MS2BP-NLS-GFP (Fig. 3C), which was very similar to the RNAscope approach (Fig. 3D). Taken together the analysis of transcriptionally active genomes in living cells supports our findings that early in infection transcription occurs from a subpopulation of imported genomes that probably require to actively remodel the viral chromatin for transcription onset. Longer observations of individual nuclei (>20min) confirmed the existence of transcribing and non-transcribing genomes with no indication of transcriptional alternation between genomes (data not shown).

**Figure 3.**
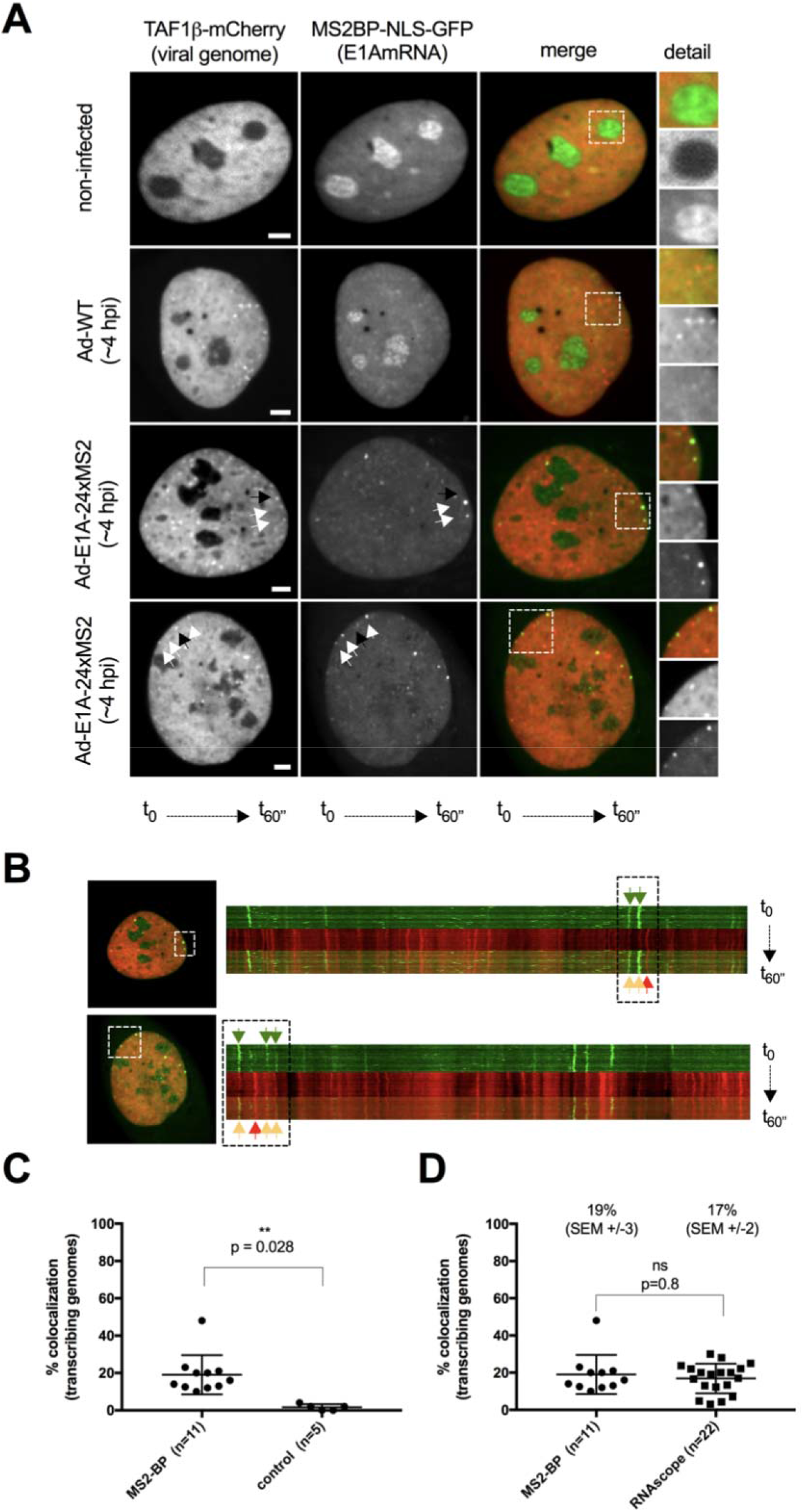
E1A transcription in living cells. **A)** Representative images showing t-projections of 60 frames (frame rate 1F/sec) imaged at ∼4 hpi. Cells express mCherry-TAF-Iβ showing viral genomes (first column, red signal in merge) and MS2BP-NLS-GFP showing MS2-repeat tagged E1AmRNAs (second column, green signal in merge) and an overlay (third column, merge) of both channels. The first row shows non-infected control cells, the second row cells infected with untagged control virus and the third and fourth row examples of cells infected with virus expressing E1AmRNA tagged with MS2-repeats. Black arrows point at non-transcribing and white arrows at transcribing genomes. Details are shown as magnifications of the boxed area. Scale bar is 10µm. **B)** Kymograph analysis of the bottom two cells shown in A and as described in the material and methods section. Cell overview to the left, kymograph separated into the individual channels and a superimposition of both channels to the right. Genome position over time appears as red vertical line, E1AmRNA signal as green vertical line. The boxed area in the overview corresponds to the boxed area in the kymograph. Green arrows point at the E1AmRNA signal, orange arrows at transcribing (double positive) and red arrow at non-transcribing (single positive) genomes. **C)** Quantitative kymograph analysis. Scatter plot representation of the percentage of MS2-BP positive genomes per cell (n = 11 cells) vs. control cells infected with viruses expressing untagged E1AmRNA (n = 5 cells) determined by kymograph analysis. **D)** Scatter plot representation comparing the percentage of E1AmRNA positive genomes per cell determined with the MS2-BP method (left, n = 11 cells) vs. the RNAscope method (right, n = 22 cells). Unpaired t-test was used for statistical analysis in C and D.

### Chromatin architecture of the invading Ad genome

Transcripts accumulated with a delay of 1 hour after nuclear entry, opening the question of whether structural changes may be required for assembly of the transcriptional machinery. To address the Ad pVII nucleoprotein organisation, we partially digested viral and cellular genomic DNA with the endo-nuclease MNase (Fig. 4A). This enzyme preferentially hydrolyses DNA in the linker region between nucleosomes and is restricted by stable protein-DNA complexes (Noll et al., 1975). Incubation of cellular chromatin with increasing concentration of MNase resulted in the appearance of a typical MNase ladder, reflecting the monomer and multimers of the nucleosomal DNA fragments (Fig. 4A). MNase concentrations were titrated such that one sample exhibited less than 20% mono-nucleosomal DNA (low digestion) and the other sample was fully hydrolysed to the mono-nucleosomal DNA (high digestion; Fig. S4A). The latter is the common condition used for MNase-seq experiments in order to study nucleosome positions (Diermeier et al., 2014; Schones et al., 2008; Valouev et al., 2011), whereas approaches using limited MNase concentrations have been used to analyse chromatin accessibility (Hu et al., 2019; Mieczkowski et al., 2016; Schwartz et al., 2018). High and low digestion conditions were used at each infection time point to isolate the mono-nucleosomal DNA band from two replicates. The nucleosomal DNA of 17 individual experiments was used for library preparation and paired-end high-throughput sequencing, yielding in total 3.8 billion paired-end reads (Table S2).

**Figure 4.**
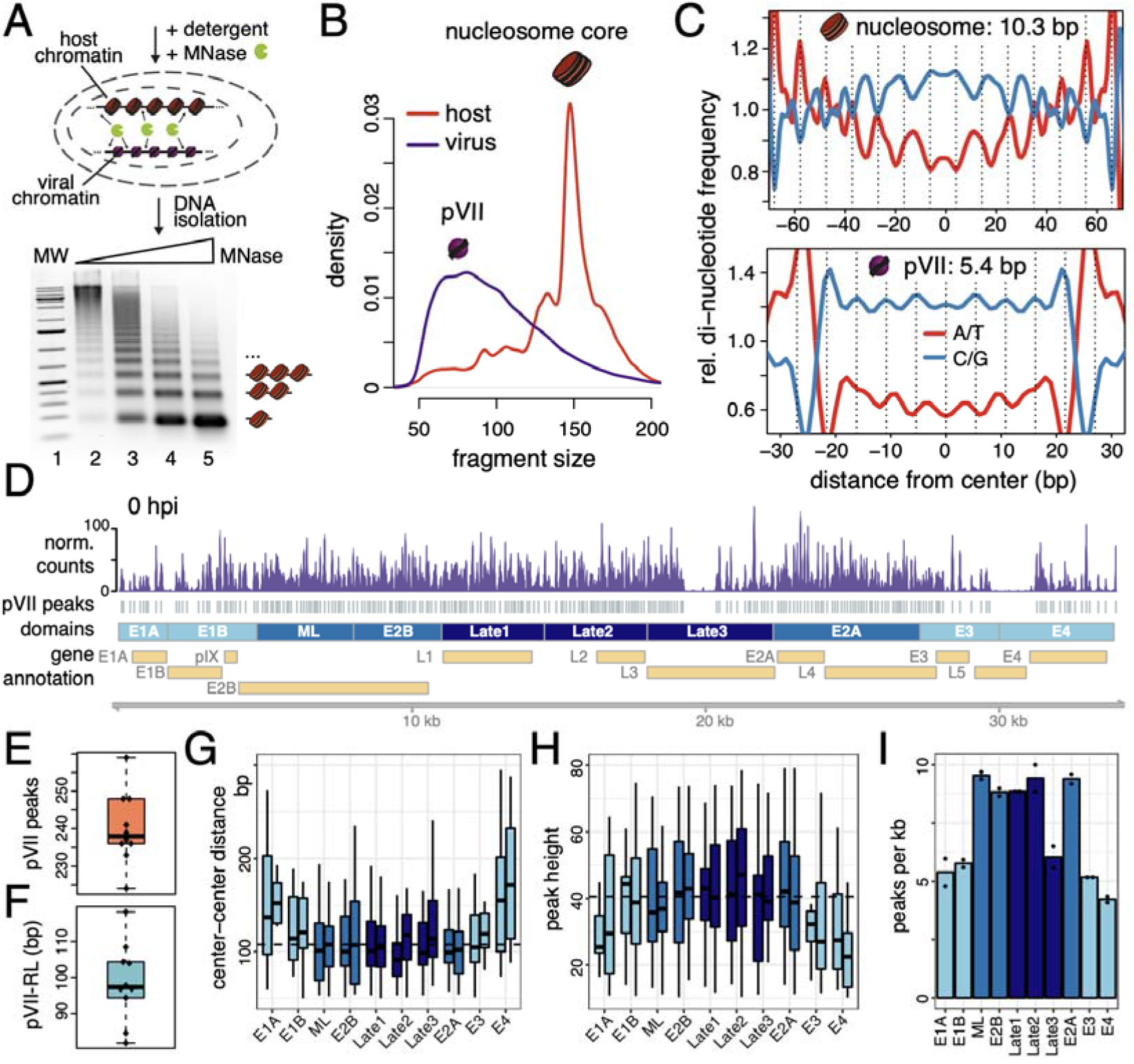
Characterization of Ad DNA packaging by pVII. **A)** Schematic view of MNase digestion of infected cells. Agarose gel shows nucleosome ladders of a MNase titration. Mono-, di- and tri-nucleosomes are indicated at the right. The mono-nucleosomal DNA fraction was isolated after MNase digestions and subjected to high-throughput sequencing. **B)** Fragment size distribution of paired-end reads either mapped to the human genome (red) or the Ad genome (purple). **C)** Average WW (A/T)- and SS (C/G)-di-nucleotide frequency of nucleosomal (upper panel) and pVII fragments (lower panel). Size-selected fragments were centred at the midpoint and the di-nucleotide frequency was calculated. Every 10.3 (nucleosome) / 5.4 (pVII) bp a dashed line highlights the periodicity of the signal. **D)** Profile illustrates Ad genome coverage by MNase-seq reads. pVII positions, which have been called with the DANPOS2 package (Chen et al., 2013), are represented by grey boxes. Partitioning of the genome into functional domains and the timing of their transcription is indicated by the color code (lightblue = early, darkblue=late). Genomic location and underlying gene annotation of HAd-C5 (NCBI accession: AY339865.1) are indicated at the bottom. **E)** Boxplot showing the number of detected pVII peaks for each sample. **F)** Boxplot showing the genome-wide pVII repeat length (pVII-RL) for each sample. **G)** and **H)** Boxplots showing the distribution of pVII center-center distances / peak heights in each domain. Individual replicates are shown side by side. **I)** Barplot showing the number of pVII peaks per kb detected in the domains. Individual replicates are shown as black dots.

DNA fragments were aligned to the human and the Ad genome, first analysing the read length distributions on both genomes (Fig. 4B). For the initial viral chromatin analysis we chose the samples at time point 0 hpi. At this stage of infection, the genome has not been released from the capsid (Fig. S1A) and therefore the MNase digestion pattern displays the nucleoprotein organization of the virus particles. When comparing the virus and host genomes digested at the high MNase concentration, we observed a clear difference in the read-length distribution on the two genomes (Fig. 4B). Whereas most DNA fragments exhibited the typical nucleosomal DNA size on the human genome (147 bp), the protected DNA fragments on the Ad genome were significantly shorter with a mean size of 60 to 70 bp. The MNase-seq approach revealed a stable adenoviral packaging unit that most probably represents pVII bound to DNA, which is the only abundant genome binding protein present prior to genome release from the capsid. Previous studies showed that nuclease digestion of Ad nucleoprotein resulted in discrete populations of protected DNA fragments, suggesting a chromatin-like configuration with sizes varying between 180 bp and 200 bp, between 30 and 90 bp, or between 30 and 60 bp depending on the conditions used (Corden et al., 1976; Daniell et al., 1981; Johnson et al., 2004). Assuming that the larger DNA fragments represent partial nuclease hydrolysis products, the 60 to 70 bp long Ad-DNA fragments mirror the previously described pVII-DNA protein complexes. Detection of these complexes (referred to as pVII peaks) allows us for the first time to analyse the occupancy and positioning of Ad pVII. We identified defined pVII peaks over the Ad genome, showing specific pVII-DNA positions on the viral DNA (Fig. 4D). The MNase profiles and peak calling were highly reproducible between biological replicates at individual time points, suggesting that specific Ad genome positions are occupied with pVII, equivalent to nucleosomes on the host genome (Fig. S4C-E). The Ad exhibits a defined nucleoprotein architecture that differs along its functional regions. Interestingly, a central region of the viral genome (Late3) and a region between the E3 and E4 genes exhibited almost no peaks (Fig. 4D).

Regions lacking pVII peaks could result either from the absence of pVII protecting the DNA from continuous MNase degradation or from tight compaction excluding DNA from MNase digestion and therefore resulting in DNA fragments of higher molecular weight. To test both possibilities we digested Ad chromatin directly within virus particles or protein-free viral DNA (gDNA, to control for MNase cleavage preferences) with a very low amount (5U) of MNase (Fig. S5A and S5B). While cleavage patterns were uniform in Ad gDNA, the highest cleavage frequencies in virus particles corresponded to the regions lacking pVII signals after infection (Late3 and between E3 and E4), suggesting the presence of “unprotected” domains in the Ad genome, which are completely hydrolyzed under the MNase digestion conditions used in infected cells.

### pVII exhibits DNA sequence dependent positioning patterns

We next asked whether intrinsic DNA sequence patterns may establish pVII peaks. Such a sequence dependent positioning code was shown for nucleosomes (Kaplan et al., 2009). Sequence specific bending of DNA reduces the energy requirements for wrapping the DNA around the histone octamer surface, revealed by characteristic di-nucleotide repeats in the nucleosomal DNA sequence at 10 bp intervals (Widom, 2001). This pattern was clearly observed in the human host genome (Fig. 4C – upper panel).

Interestingly, we also observed a specific di-nucleotide pattern when computing the di-nucleotide frequency of the mid-point centered pVII peak fragments, suggesting that a specific DNA code underlies pVII positioning on the Ad genome (Fig. 4C – lower panel). However, to our surprise, a periodic pattern of 5.4 bp was obtained for WW-(where W is A or T) and SS-dinucleotides (where S is G or C), much shorter than the nucleosomal DNA repeat pattern, implying a distinct packaging mode. Such a repeat pattern indicates a spring-like winding of the DNA around pVII. Even without knowledge of the exact DNA-pVII topology, the sequence pattern and the specific binding sites of pVII on DNA clearly suggest the organisation of the Ad genome into a specific nucleoprotein-structure that may be essential for its function.

On average, we detected 238 pVII peaks covering the Ad genome, which was consistent throughout all timepoints (Fig. 4E). Next, we quantified the mean pVII-DNA particle repeat length (pVII-RL), the distances between individual pVII peaks, by measuring the center-center distance of adjacent pVII peaks in each sample (Fig. 4F and S4B). The median pVII-RL measures 97.3 bp, suggesting that we isolated defined pVII-DNA particles spaced by accessible DNA linkers of 20 to 40 bp in length.

To address the presence of functionally distinct regions, we systematically subdivided the Ad genome into 10 domains of similar size, named after the included gene and classified by the timing of expression (color code: early to late = light to dark blue) (Fig. 4D). Plotting the pVII-RL showed a clear correlation of the repeat length with expression timing (Fig. 4G). Increased repeat lengths in the regions of the early genes may serve to increase genome accessibility and to enable the binding of regulatory factors. Strengthening this finding, the pVII occupancy and stability, reflected by the peak height, showed an anti-correlation with early timing in gene expression (Fig. 4H). In general, less pVII peaks (Fig. 4I) with larger repeat lengths and increased instability were found at early expressed domains (Fig S4D-E) indicating an open chromatin conformation, which could be confirmed with limited MNase digestions of virus particles (Fig. S5D).

Our high-resolution analysis of Ad genome packaging provides, for the first time, information on pVII-DNA peak positioning and occupancy. The analysis revealed a defined viral nucleoprotein architecture, with an open configuration at early expressed genes potentially enabling immediate access of the host cell transcription machinery.

### Dynamic changes of viral genome accessibility over time

One parameter determining viral genome accessibility and activity is the specific occupation with pVII, another parameter is the higher order structure of the Ad-chromatin and its global accessibility. Assessing the nucleoprotein structure with high MNase concentrations is providing information about stable protein particles occupying the DNA (Schwartz et al., 2018). In contrast, limited MNase concentrations, in analogy to the analysis of regulatory sites by DNase I hypersensitivity mapping (Längst et al., 1997), can reveal accessible genomic regions and regulatory elements (Schwartz et al., 2018).

Only a fraction of about 20% of the viral genomes in the nucleus are transcriptionally active (Fig. 2E and 3C), explaining why we observed only little variation between the different time points treated with high MNase concentrations (Fig. 5A). In contrast, at low MNase concentration (Fig. 5B and Fig. 5C) clear changes over infection time are detected, signifying gross structural changes of the viral chromatin during early infection. We hypothesize, that the active viruses undergo de-compaction that can be detected by limited MNase hydrolysis. When monitoring the pVII peak positioning at 0 hpi, the low MNase signal was well correlated with the high MNase signal (R = 0.91), essentially showing the same pVII peak positions in both conditions (Fig. S6A, B). However, when inspecting the viral chromatin maps at later time points, we detected specific structural changes, with changes in peak intensities and the appearance of novel peaks in the low MNase signal (Fig. 5C). To identify structural changes over time we used a linear regression approach. The average signal intensity at each pVII position was quantified at each time point and fitted by linear regression. The slope of the regression line was used as a score for chromatin accessibility changes (accessibility score; Fig. 5C). Genomic regions with a positive accessibility score are opening up, being more MNase accessible, whereas negative values correlate with DNA remaining condensed. A similar strategy was used in previous studies to measure the local chromatin accessibility (Chereji et al., 2019; Mieczkowski et al., 2016; Mueller et al., 2017). The increase of Ad chromatin accessibility is correlating with viral transcriptional activity, with Ad chromatin remodeling and de-condensation at the early gene regions (Fig. 5D). Interestingly, chromatin opening at E1A is not as pronounced as at other early genes. As this is the first gene to be transcribed, we suggest that this region already exists in an open chromatin configuration, correlating with the pre-existing low pVII peak density, and therefore no additional opening is observed over time.

**Figure 5.**
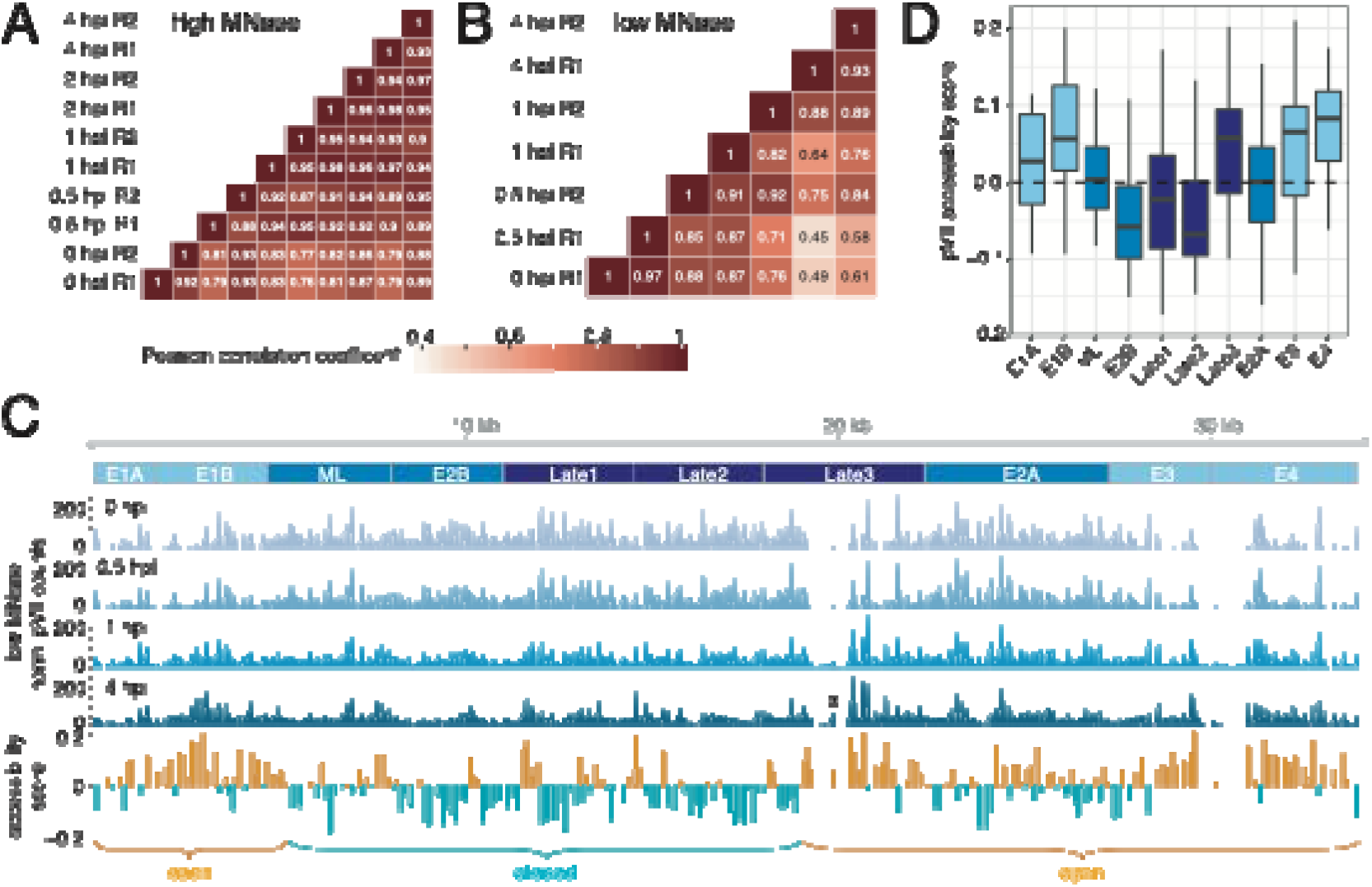
Early decondensation of Ad chromatin. **A)** and **B)** Sample-wise Pearson correlations of **A)** high MNase or **B)** low MNase profiles. **C)** Profile illustrates Ad genome coverage by low MNase-seq fragments. The average of two replicates is shown, except at timepoint 0 hpi where only one replicate was available. The accessibility score was calculated as the slope of a linear regression fit of the average signal intensity across each timepoint. The score was assessed for each peak. Yellow bars depict increased accessibility and blue bar show peaks with reduced DNA accessibility. Domains of open and closed Ad chromatin are shown below. A new accessibility peak arising during infection in the Late3 region is marked by an asterisk. **D)** Barplot showing the average accessibility score in each domain.

The Late3 domain also exhibited a positive accessibility score, suggesting Ad chromatin de-condensation at this site (Fig. 5C). Additionally, in this domain we observe a discrete peak appearing at low MNase concentrations at 4 hpi (Fig. 5C shaded region and peak marked by asterik; Fig. S6C). The occurrence of this local nuclease hypersensitive site suggests the binding of regulatory factors (Chereji et al., 2017; Kent et al., 2011; Mieczkowski et al., 2016). Transcription factor motif analysis revealed potential binding sites for EN1, CREM, C/EBP-alpha, and C/EBP-beta in the center of the peak (Fig. S6C). Interestingly, the transcripts of EN1, CREM, and C/EBP-beta in the host cell were upregulated up to twofold upon viral infection (Fig. S6D).

In summary, the time-resolved chromatin accessibility analysis shows, that in addition to the predefined pVII binding sites on the genome, Ad chromatin compaction specifically changes over time. Chromatin opening of the early gene regions starts at 0.5 hpi, prior to transcriptional activation. We suggest that changes in early gene regions are a pre-requisite for gene activation process.

### Dynamic changes of Ad genome packaging

Previous studies showed that histones are deposited onto the Ad genome (Giberson et al., 2018; Komatsu and Nagata, 2012; Komatsu et al., 2011). However, at which position and whether these are functional nucleosomes is not known.

Analysing the MNase-seq data of the time points 1, 2 and 4 hpi revealed the appearance of 147 bp long, protected DNA fragments at 2 and 4 hpi, after nuclear import of the viruses and prior to early gene activation (Fig. 6A). The nucleosome sized peaks increased in intensity, within the background of the more abundant pVII protected DNA fragments, suggesting co-occupation of the viral genome by pVII and nucleosomes. To quantify the fraction of nucleosomes on the viral genome, we simulated the peak sizes by computationally adding nucleosomal reads to the pVII fragment data of time point 1 hpi (Fig. S7A). The analysis suggested that at 2 hpi 8% and at 4 hpi 14% of the viral genomes are occupied with nucleosomes.

**Figure 6.**
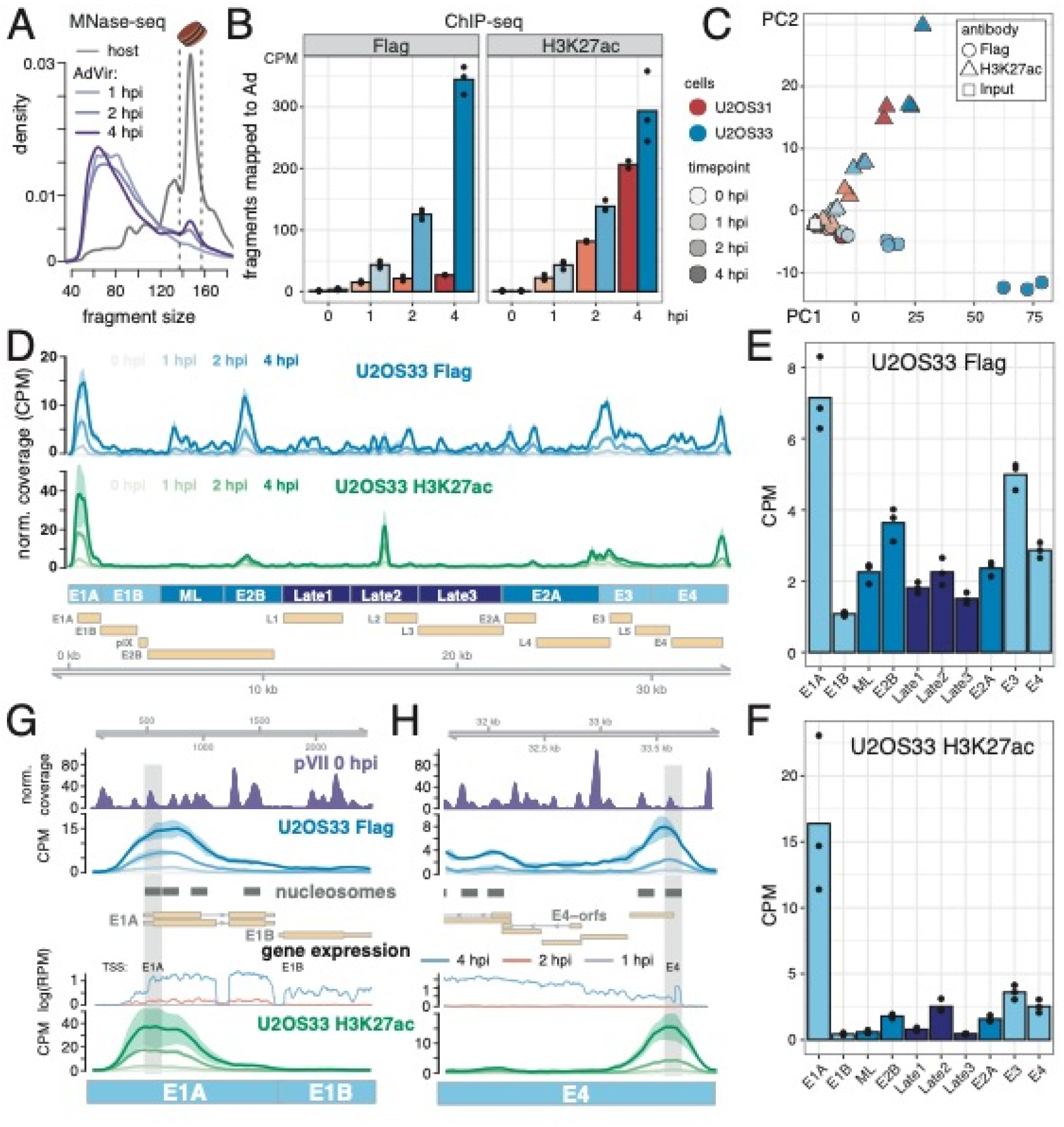
Nucleosome assembly onto the Ad genome. **A)** Fragment size distribution of paired-end reads from high MNase experiments (see Fig. 4) either mapping to the human genome (grey) or the Ad genome (shades of purple). Nucleosome sized fragments are highlighted between two dashed lines at 137 bp and 157 bp. **B)** Quantification of ChIP-seq fragments mapping to the Ad genome over time. Flag/H3K27ac immunoprecipitation (IP) was conducted in U2OS cells expressing either Flag-tagged H3.1 (U2OS31, blue shades) or H3.3 (U2OS33, red shades). The barplots show the average of 2-3 replicates (black dots) of each time point and condition. The antibody used in the IP is indicated at the top. **C)** Principal component analysis (PCA) of ChIP-seq samples. **D)** H3.3 and H3K27ac profiles along the Ad genome. Normalized coverage of Flag (blue shades) / H3K27ac (green shades) ChIP-seq experiments in U2OS33. The average signal of 3 replicates is shown and the 95% confidence interval is indicated. CPM = counts per million mapped fragments **E)** Average coverage of Flag-tagged H3.3 or **F)** H3K27ac in specific Ad domains. **G)** and **H)** Genome browser tracks showing pVII occupancy at 0 hpi (upper panel), the H3.3 (blue shades) / H3K27ac (green shades) accumulation over time, predicted sites of nucleosome assembly (grey boxes) using nucleosome-sized fragments (137-157 bp) from 4 hpi MNase-seq samples and expression levels of early genes (middle panel).

To verify this indirect prediction of nucleosomal occupancy on the Ad genome, we generated U2OS cell lines, stably expressing FLAG-tagged histone H3.1 (U2OS31) and the histone variant H3.3 (U2OS33) (Fig. S7B, C). Stable cell lines were infected with Ad and ChIP-seq experiments were performed at 0, 1, 2, and 4 hpi, using either FLAG-or H3K27ac-antibody. At 0 hpi we barely detected any ChIP-seq reads mapping to the Ad genome, showing the high specificity of our ChIP-seq approach (Fig. 6B). The number of precipitated Ad DNA fragments increased with infection time for histone H3.3 (FLAG IP: U2OS33), in contrast to the canonical histone H3.1 DNA fragments (FLAG IP: U2O31) (Fig. 6B). The low number of histone H3.1 DNA fragments clustered with the samples taken at 0 hpi on PCA and hierarchical clustering analysis (Fig. 6C, Fig. S7D), and did not show any specific sites of enrichment on the Ad genome (Fig. S7E, upper panel), suggesting no specific recruitment of histone H3.1. However, we detected specific H3.3 binding sites on the Ad genome, clearly showing that this histone variant is exclusively deposited onto incoming viral genomes (Fig. 6D). The H3.3 ChIP-Signal was most pronounced at early transcribed gene promoters, including E1A, E3, and E4, but not E1B. Nevertheless, H3.3 also accumulated at additional loci throughout the Ad genome, like the E2B and Late2 regions (Fig 6D, E).

Next, we performed H3K27ac ChIP-seq experiments, showing efficient histone acetylation, without any detectable delay after histone deposition on the Ad genome. Interestingly, we observed histone acetylation at specific sites on viral genomes, representing a subset of all H3.3 binding sites, marking these as active regions (Fig. 6B, D). As the H3K27ac antibody does not discriminate between H3.1 and H3.3, the results obtained in U2OS31 and U2OS33 are very similar (Fig. 6B, C and Fig. S7D). The U2OS33 cell line exhibited only slightly higher H3K27ac levels due to additional expression and incorporation of histone H3.3. Histone acetylation was identified at all early gene promoters, except E1B, with the strongest signal at E1A, which was already detectable 1 hpi (Fig. 6D, F). Additionally, we observed pronounced K27 acetylation at two central regions covering the E2B or Late2 gene.

We expected that sites marked by H3K27ac would promote chromatin de-condensation. However, neither H3.3 deposition nor H3K27 acetylation did correlate with chromatin accessibility changes (R^2^ < 0.1, Fig. S8A, B). A closer inspection showed that acetylation at the E1A and E4 genes were associated with open chromatin (Fig. S8C, F), whereas the acetylated Late2 and E2B sites in the genome centre remain condensed (Fig. S8D, E).

In summary, we suggest that the histone variant H3.3 is preferentially deposited onto viral genomes and a subset of binding sites get acetylated at K27. H3.3 acetylation is detected preferentially at early gene promoters, being nucleosomal and acetylated prior to transcription initiation.

### Early genes gain a positioned +1 nucleosome

The sonicated DNA in the ChIP experiments is not short enough to reveal the exact sites of nucleosome assembly. Therefore, we combined the ChIP-seq data with the MNase-seq derived nucleosomal DNA fragments (size between 137-157 bp; time point 4 hpi), allowing us to reveal the exact positions of the nucleosome core particle. A cut-off of 2 CPM was used to define the genomic sites of preferential nucleosome assembly from U2OS2 FLAG IPs (Fig. S8G, blue boxes) and the exact nucleosome positions were determined using MNase size selected fragments (Fig. S8G, golden boxes). The nucleosome profile was distinct from the pVII landscape and nucleosome calling was reproducible between replicates.

Nucleosome assembly occurs in the genomic regions associated with the early genes, exhibiting a lower pVII peak occupancy and increased pVII peak repeat-length. An analysis of the incoming nucleosome positions with respect to pVII peak occupancy revealed that nucleosomes directly formed over the central positions of the pVII-nucleoprotein particles (Fig. S8H) suggesting a specific mechanism to replace pVII.

A detailed analysis of the nucleosome assembly sites with respect to the early genes revealed that nucleosomes occupy discrete positions. Remarkably nucleosome positions coincide with the 5’-end of the synthesized E1A, E4, E3, and E2A RNA, correlating with the highly positioned +1 nucleosome found in active human genes (Fig. 6G, H and Fig. S8I) (Schones et al., 2008). Additional nucleosomes are detected in the coding region of the early genes. The +1 nucleosome marks the transcription start sites of genes and flanks the regulatory sequences of the promoters, residing in the upstream DNA linker region.

The time-resolved chromatin analysis shows, that the predefined pVII chromatin organisation is remodeled, exchanging pVII for histone H3.3. Histone H3.3 deposition and acetylation occur mainly at the +1 region of early genes and happens prior to transcription onset. Our results imply the requirement of specific nucleosome assembly for transcription initiation at early genes.

## DISCUSSION

### The adenovirus exhibits a defined nucleoprotein architecture

Early studies suggested that pVII binds adenoviral DNA at a 1:1 ratio like histones and protects DNA from endonuclease cleavage (Lischwe and Sung, 1977; Sato and Hosokawa, 1984). However, no defined pVII-DNA particle has been identified on functional Ad genomes yet, as the protected DNA had a variable size range of 30 to 250 bp depending on experimental conditions and the degree of endonuclease digestion (Corden et al., 1976; Daniell et al., 1981; Mirza and Weber, 1982). In addition, it is still not known whether pVII binds to specific sites on the viral genome, creating a defined viral nucleoprotein architecture, like the cellular nucleosomes. Here, our MNase-seq approach digesting Ad together with the host chromatin at different time points post infection, revealed the 147 bp long DNA fragments of the host nucleosomes and identified a shorter, protected DNA species with a mean size of about 60 to 70 bp originating from the viral sequences. With pVII being the major DNA binding protein of the adenovirus, and the protected fragment species being present prior to genome release from the capsid, the observed DNA fragments are very likely protected by their association with pVII. The viral MNase digestion pattern is not as regular as the nucleosomal DNA fragments, suggesting heterogeneity in pVII-DNA complexes, as observed in the early studies. Such a digestion pattern could also be the result of a discrete pVII core complex (60 bp) that tightly associates with neighbouring pVII complexes, partially protecting the interconnecting linker DNA from endonuclease cleavage. Such structures were indeed visible in early electron microscopic studies (Mirza and Weber, 1982).

### pVII organizes the viral DNA into a specific nucleoprotein structure

Annotation of the putative pVII protected DNA fragments to the viral genome revealed for the first time a specific positioning of pVII peaks on the genome. We show that the Ad genome is covered by an average of 238 defined pVII peaks located at defined positions that are spaced by DNA linkers of about 20 to 40 bp. Precise pVII peak positions suggest a highly organized and defined nucleoprotein architecture shared by all virus particles. In host cells, DNA structure determines histone binding preferences, as shown by the DNA sequence dependent nucleosome positioning code (Kaplan et al., 2009). DNA bending allows wrapping of the DNA around a protein core, requiring site specific SS and WW di-nucleotide enrichments. SS and WW repeats differ in the wideness of the DNA major/minor groove thereby bending the DNA (Kaplan et al., 2009). If the sequence dependent DNA shape follows the protein surface path it increases binding affinity and thereby determines binding site preferences (Widom, 2001). Whereas the nucleosomal di-nucleotide repeats are spaced by 10.3 bp, pVII exhibits a much shorter repeat length of 5.4 bp. Nucleosomes and pVII-DNA complexes are organized differently, and we suggest that the DNA associates in form of a spiral around the pVII core (Fig. 4C).

Our results suggest that pVII folds Ad genomes into an organized higher order structure, driven by specific binding sites pre-determined by the Ad DNA sequence. A recent report showed that pVII is not required for genome packaging into virions, albeit the lack of pVII resulted in non-infectious Ad (Ostapchuk et al., 2017). The specific organization of the viral genome with pVII may be functionally linked with endosomal escape, in addition to potential gene regulatory effects. However, as endosomal escape of the Ad lacking pVII is blocked and genomes are not delivered to the nucleus, the effect of pVII absence on gene expression and the specific exchange of positioned pVII peaks by nucleosomes cannot be concluded. Our observation that pVII peaks are reproducible across biological replicates strongly indicates a specific positioning code. Protein VII peak densities and the changes in pVII occupancy upon nuclear delivery (see below) correlate with the onset of the transcriptional program and cellular factors, such as TAF-Iβ. TAF-Iβ was shown to positively influence viral transcription, being recruited to the incoming genomes via pVII (Haruki et al., 2003; Matsumoto et al., 1995). One possible approach to validate our findings of an underlying sequence code for pVII positioning may be the rational (sequence-)design of therapeutic DNAs packed into adenoviral vectors to achieve high and sustained transgene expression. *In vitro* assembly of pVII-DNA complexes showed that pVII binds to DNA regardless of the DNA sequence (Avgousti et al., 2016; Komatsu et al., 2011). However, it was not addressed, whether sequence motifs are preferred by pVII, like it was observed in nucleosome assemblies. Specific assembly factors and ATP dependent remodeling enzymes may be required to adjust the precise positioning of pVII on DNA, in addition to the DNA assembly code. Like nucleosomes, pVII does not bind as isolated protein to DNA but is forming homotypic complexes. The number of pVII proteins per Ad genome was estimated to be about 500 copies (Benevento et al., 2014) and early biochemical studies suggested the occurrence of pVII dimers (Chatterjee et al., 1985; Everitt et al., 1975). Taking into account the median number of 238 pVII positions we identified on the Ad genome (plus 19 on average for the 1871 bp deleted E3 region) our data would be compatible with two pVII molecules forming dimers at 238 discrete sites. This number would also come close to the ∼200 “beads-on-string” estimated for purified viral cores (Vayda et al., 1983).

### Viral chromatin dynamics drives early gene expression

Here we show that nuclear delivery of viral genomes takes about 30 min but requires an additional hour before viral transcripts appear. In agreement with earlier observations, full activation of immediate early gene expression is reached after 4 h (reviewed in (Pied and Wodrich, 2019)). Correlating gene expression with our differential MNase digestion conditions, we show that early genes exhibit a reduced density of MNase protected regions that most probably represent pVII peaks, and increased pVII peak repeat lengths. Direct detection of genomes and of the E1A mRNA showed that genomes imported to the nucleus require about two hours to activate gene expression and to synthesize the mRNA (i.e. E1A transcription). In this lag phase pVII is selectively replaced by nucleosomes at early genes, preceding gene activation. ChIP-seq experiments revealed that the histone variant H3.3 is incorporated at many specific genomic locations on the Ad genome, including the early gene regions, shortly after nuclear import. A subset of these nucleosomal sites, corresponding mainly to the early gene positions, are acetylated at H3K27 prior to transcriptional activation.

We identified that 80% of the imported Ad genomes are transcriptionally inactive, but the remaining 20% do actively rearrange the chromatin structure to allow transcription factor and RNA Polymerase binding. Once entering the nucleus, we showed that the chromatin structure of active viral genomes decondensed and viral DNA becomes more accessible. A delay for E1A gene expression and heterogeneity of transcriptional activity for imported genomes was recently reported supporting our findings (Suomalainen et al., 2020). The limited activation of incoming genomes and subsequent de-compaction was also observed during herpes simplex virus type 1 (HSV-1) infections showing common features of nuclear replicating DNA viruses (Hu et al., 2019; Sekine et al., 2017). However, in the case of HSV-1 gene activation, it was suggested that the accessibility over the complete viral genome increased (Hu et al., 2019), whereas active Ad genomes selectively de-condense the early gene regions and a specific domain in the Late3 gene. The overlap of open viral chromatin with histone H3K27 acetylation and early transcription, suggests an organization that allows the binding of regulatory factors for gene activation. In contrast, for the Late3 region no functionality in gene expression control has been assigned yet. Our observation that increased DNA accessibility overlaps with binding motifs for transcriptional regulators may indicate a role as a transcriptional enhancer.

### pVII is exchanged for nucleosomes at early genes

Genome de-compaction correlates with low pVII occupancy and coincided with a site-specific exchange of pVII protein for histone octamers. The characteristic nucleosomal fragment length of 147 bp, observed in MNase-seq, confirms the release of pVII from the viral genome, as pVII-nucleosome complexes would result in longer fragments after MNase digestion as previously shown (Avgousti et al., 2016). The ChIP-seq experiments clearly show an overlap of the 147 bp long protected DNA fragments and the location of the histone variant H3.3. Furthermore, the newly assembled nucleosome is located on a previous position of a pVII-DNA complex. Once pVII is set free, it is able to sequester HMGB1 and modulate the cellular chromatin structure, thereby downregulating the immune response of the host cell (Avgousti et al., 2016). Protein VII molecules were exchanged for positioned nucleosomes at the +1 site of actively transcribed genes (i.e. the E1A and E4 gene, Fig. 6G and 6H). The calculated and experimentally validated number of nucleosomes and their positions on DNA, as determined by the 147bp long DNA fragments and the ChIP-seq experiments, corresponds to the number of ∼ 20% transcribing genomes, exhibiting single nucleosomes positioned at the +1 position of early genes. As nucleosome exchange is commencing at 0.5 hpi and is completed at 2 hpi when transcription activation is detectable by mature transcript accumulation, we suggest that nucleosome positioning is a prerequisite for gene activation. At this point we can only speculate why only a subpopulation of genomes is transcriptionally active. Reasons could be that the few genomes undergoing nucleosome assembly and active transcription produce the replication enzymes, whereas the bulk of genomes enters replication without activation as an elegant way to avoid repeated chromatinization. Alternatively, cellular defense mechanisms and/or the cell cycle could influence transcription levels (Suomalainen et al., 2020).

Interestingly, pVII removal and nucleosome placement at early genes preceded transcription, suggesting that pVII removal is driving transcription and not vice versa, as previously reported (Chen et al., 2007). This observation is in agreement with recent pVII-ChIP experiments showing transcription and replication independent pVII removal in early infection (Giberson et al., 2018; Komatsu and Nagata, 2012; Komatsu et al., 2011).

Future work has to reveal which machinery drives the pVII nucleosome exchange and determines specificity. Generally, active promoters are free of nucleosomes (nucleosome depleted region, NDR) and the transcription start site is typically located upstream of a well-positioned ‘+1’ nucleosome influencing recruitment of the transcriptional machinery and initiation of transcription (Jiang and Pugh, 2009; Kubik et al., 2015). In the case of the Ad genome, our data show that the E1A promoter region is already pVII free and exists in an accessible nucleoprotein conformation when entering the cell, potentially allowing fast assembly of the +1 nucleosome and enabling immediate gene activation. This organization might thus be defined in the producer cell during genome packaging, which occurs in a sequential packaging sequence driven manner (Hammarskjöld and Winberg, 1980; Hearing et al., 1987). In contrast, other early regions were actively de-compacted with nuclear entry of the virus, correlating with slightly delayed transcriptional onset. This effect may be driven by pVII interaction with transcription promoting factors like TAF-Iβ and E1A (Haruki et al., 2006; Johnson et al., 2004). Besides pVII is post-translationally poly(ADP)-ribosylated or acetylated in infected cells, which might further promote chromatin relaxation (Avgousti et al., 2016; Déry et al., 1986).

Taken together our analysis provides unprecedented insight into the adenoviral nucleoprotein structure and its dynamic reorganization during the onset of viral gene expression. We reveal genome sequence, pVII-DNA occupancy and nucleosome positioning as driving forces for transcriptional activation. Reproducing such organization in adenoviral vectors could result in efficient and sustained transgene expression.

## EXPERIMENTAL PROCEDURES

### RNA library preparation and sequencing

Total cellular RNA was isolated from infected H1299 cells at 0, 0.5, 1, 2, 4 hpi using TRI Reagent (Molecular Research Center, Inc.). Two time series sets of total RNA were prepared from two independent infections. Library preparation, comprising rRNA depletion, polyA enrichment, fragmentation to ∼270 nucleotide length, reverse transcription to cDNA using random hexamer primers and adapter ligation, was accomplished using TruSeq RNA Sample Preparation Kit v2 (Illumina) according to the manufacturers protocol by EMBL GeneCore facility in Heidelberg. Libraries were sequenced on Illumina HiSeq2000 platform resulting in 18-75 Mio 50 bp single-end reads per sample.

### Differential MNase-seq, library preparation and sequencing

H1299 cells infected with Ad were permeabilized with the detergent IGEPAL-CA630 and digested at 0, 0.5, 1, 2 and 4 hpi with 100U/600U of MNase for 4/5 min at 37°, denoted as ‘low’/’high’ MNase. Mnase treatment was performed as described in (Schwartz et al., 2018). After fragmented DNA has been separated on 1.3% agarose gels, mono-nucleosomal fractions were isolated. Paired-end sequencing on Illumina HiSeq2000 platform (EMBL GeneCore facility, Heidelberg), yielding about 220 million read pairs of 50 bp length per sample, was accomplished following library preparation using NEBNext DNA library prep Master Mix Kit (New England Bioloabs)

### ChIP-seq, library preparation and sequencing

Chromatin immunoprecipitation (ChIP) was performed as described previously with slight modifications (Minderjahn et al., 2020). Chromatin of cell lines encoding Flag-tagged H3.1 or H3.3 (U2OS31 and U2OS33, respectively) was harvested at 0, 0.5, 1, 2 and 4 hpi in triplicates of independent Ad infections. Briefly, for anti-H3K27ac and anti-Flag ChIP-seq, cells were crosslinked with 1% formaldehyde for 10 min at room temperature and the reaction was quenched with glycine at a final concentration of 0.125 M. Chromatin was sheared using ultrasonication (Covaris S2). A total of 2.5 µg of antibody against H3K27ac (Abcam, ab4729), or FLAG (M2, Sigma Aldrich, F3165), was bound to 20 µl pre-washed Dynabeads Protein A (Thermo Fisher Scientific) in a total volume of 200 µl PBS containing 0.02% Tween-20 for 1h at room temperature rotating at 6 rpm. The buffer was removed on a magnet and antibody-coupled beads were resuspended in 120µl dilution buffer (20mM Tris pH7.4, 2mM EDTA, 100mM NaCl, 0.5% Triton x-100) containing proteinase inhibitors (cOmplete™, EDTA-free Protease Inhibitor Cocktail, 50x, Sigma). 80µl sonicated chromatin was added to the antibody-coupled Dynabeads (sonicated chromatin of approx. 1.7×10^6^ cells) and incubated for 3h at room temperature rotating at 6 rpm. Beads were washed on a magnet and chromatin was eluted. After crosslink reversal, RNase A and proteinase K treatment, DNA was extracted with the Monarch PCR & DNA Cleanup kit (NEB). Sequencing libraries were prepared with the NEBNext Ultra II DNA Library Prep Kit for Illumina (NEB) according to manufacturer’s instructions. The quality of dsDNA libraries was analyzed using the High Sensitivity D1000 ScreenTape Kit (Agilent) and concentrations were assessed with the Qubit dsDNA HS Kit (Thermo Fisher Scientific). Libraries were sequenced paired end on a NextSeq550 (Illumina).

### Viruses and virus production

Bacterial artificial chromosome (BAC) mutagenesis was used to generate recombinant adenoviruses following the protocol described in (Ruzsics et al., 2014) and (Martinez et al., 2015). Briefly, bp 1-3513 of the HAdV-C5 genome including the left inverted terminal repeat (ITR) and the whole E1 region was replaced by an FRT (Flippase Recognition Target) site and maintained as BAC (termed B12) to perform lambda red recombination essentially as described. Plasmid pO6-Ad5-E1-FRT (encoding the left ITR and the complete E1 region followed by an FRT recombination site) was used to clone 24 copies of the MS2 binding site repeats into the 3’UTR of the E1A gene. Subsequently flp-recombination was used (Ruzsics et al., 2014) to recombine plasmids pO6-Ad5-E1-FRT-MS2 and BAC B12 to reconstitute BxAd5-E1-MS2 a BAC encoding the whole AdV genome including the E1A gene tagged with 24 copies of the MS2 binding site. To generate the virus the BxAd5-E1-MS2 BAC was digested by *PacI* restriction enzyme to release the viral genome and transfected into HEK293 cells to reconstitute the recombinant viruses. Two rounds of plaque purification were performed followed by large scale production in HEK293 cells and purification through double CsCl_2_ gradients banding. Replicative E3-deleted HAd-C5 virus or E1/E3-deleted HAd-C5 GFP expressing control virus were essentially produced in the same way. Purified viruses were extensively dialyzed against PBS/10% glycerol and stored in aliquots at -80°C and quantified as physical particles per µL following the method described in (Mittereder et al., 1996).

### Cell culture and Infection

U2OS (ATCC #HTB-96) and H1299 (ATCC #CRL-5803) and HEK293 (kindly provided by G. Nemerow, TSRI, La Jolla USA) cells were maintained at 37°C and 5% CO_2_ in Dulbecco’s modified Eagle’s medium (DMEM)-Glutamax supplemented with 10% fetal calf serum (FCS) and 1% of Penicillin/Streptavidin. U2OS cells stably expressing mCherry-TAF-Iβ (Komatsu et al., 2016) were transduced with lentiviruses expressing MS2BP-NLS-GFP and double positive cells were sorted by fluorescence activated cell sorting (FACS). The resulting cell population was maintained without further selection. For control experiments using cytosolic MS2BP (Fig. S6) stable mCherry-TAF-Iβ cells were transfected with an expression vector for MS2BP-NES-GFP. For RNAscope and immunofluorescene (IF) analysis H1299 cells were grown on 15mm glass coverslips. Subsequently cells were infected with 3000 physical particles/cell of replicative E3-deleted HAd-C5 virus or E1/E3-deleted HAd-C5 GFP expressing control virus, washed once with PBS and fixed at different time points with paraformaldehyde 4%/PBS for 10min at room temperature. Fixed cells were washed 3 times with PBS and slowly dehydrated in 50%/70%/100% ethanol for 5 min each and stored at - 20°C in 100% ethanol. For the RNAscope protocol cells were rehydrated for 2 min in 70%/50% ethanol and equilibrated for 10 min in 1x PBS.

### Preparation of stable cell lines expression FLAG tagged H3.1 and H3.3

U2OS were seeded in 6-well plates (250 000 cells/well) and transfected with 2 µg of pcDNA3-Histone H3.1-FLAG or pcDNA3-Histone H3.3-FLAG (kindly provided by K. Nagata, Tsukuba University, Japan) using Lipofectamine 2000 (Invitrogen). Two days after transfection, cells were selected with 3 mg/ml G418 for 13 days. Resistant cells were cloned by limiting dilution and the uniform expression of the tagged histones was monitored by immunofluorescence and western-blot on individual clones.

### RNAscope

RNAscope experiment were performed using reagents and protocols from the RNAscope® Multiplex Fluorescent Assay essentially as indicated by the supplier (ACD Bio). Briefly, cells were incubated with Protease III (freshly diluted 1:30 with PBS) for 15 min at room temperature (RT) and washed 3 times with PBS. Hybridisation with the E1A RNAscope probe was performed for 2h at 40° in a humidified chamber. Hybridized cells were washed twice for 2 min and hybridized with signal amplifier probes (final amplifier Amp 4 AltB-FL) following the manufacturer’s instructions. Hybridized cells were washed twice in PBS followed by incubation in IF buffer (PBS/10% FCS/0.1% Saponin) for 15 min at room temperature to saturate unspecific binding sites followed by IF analysis. Primary antibodies (mouse anti-pVII) were diluted in IF buffer and incubated with cells for 1h at RT. Antibody was removed by washing in PBS followed by incubation with secondary antibodies (mouse Alexa 647) diluted in IF buffer for 1h at RT. Cells were washed with PBS, ddH_2_O and absolute ethanol, air-dried and mounted in DAKO mixed with DAPI (1:1000).

### Live-cell imaging and image analysis

For live-cell imaging, U2OS cells stably expressing mCherry-TAF-Iβ and MS2BP-NLS-GFP were seeded in ibidi μ-slide VI^0.4^ (Ibidi). The next day, the medium in the ibidi μ-slide was replaced by imaging medium (FluoBrite DMEM supplemented with Prolong antifade reagent, both Life technologies) containing 3000 physical particles/cell of replicative BxAd5-E1-MS2 for 30min followed by washing with imaging medium to remove the inoculum. Infection and subsequent cultivation were performed at 37°C. Cells were imaged using a Leica spinning-disk microscopy system (x100 objective) equipped with an environmental chamber at a frame rate of 1 frame per second using MetaMorph software. To generate kymographs, movies were processed using the ImageJ KymographBuilder plugin (https://imagej.net/KymographBuilder_:_Yet_Another_Kymograph_Fiji_plugin) on confocal sections. Line width was adapted to cover all viral genomes throughout the length of the movie and double positive genomes were scored as transcribing genomes. Mounted cells for IF and RNAscope analysis were imaged using a SP8 confocal microscope (Leica) at the Bordeaux Imaging Center (BIC) platform. Stacks (7x) were taken every 0,3µm using a pinhole of 1 at a pixel size of 85 nm. Quantifications were done using an adapted semi-automated macro in ImageJ (provided upon request). Briefly, channels were split and stacks were Z-projected and the cell periphery was outlined manually. A threshold was applied to every channel and single objects or colocalized objects exceeding a defined area of pixels were counted. Statistical analysis was done using one-way ANOVA multicomparison test.

## QUANTIFICATION AND STATISTICAL ANALYSIS

### Virus annotation

The reference genome sequence and gene annotation of Human Adenovirus C serotype 5 (accession: AY339865.1) was downloaded from NCBI. The annotation was adapted to the used Adenovirus strain by excising nucleotides 28,593-30,464 from the full version (modified virus reference genome and gene annotation available at the github project page).

### RNA-seq analysis

STAR (Dobin et al., 2013) was used to align the RNA-seq reads simultaneously to the human and the adenoviral genome. Therefore, a STAR index was generated based on the hg19 reference, the modified adenoviral genome and gene annotation (see section *Virus annotation* for details) and the GENCODE human transcript annotation version 25. Read mapping was conducted using following parameters: --outSAMmultNmax 1 --outMultimapperOrder Random --outFilterType BySJout --outFilterIntronMotifs RemoveNoncanonical -- outSAMstrandField intronMotif --outFilterMultimapNmax 20 --alignSJoverhangMin 8 -- alignSJDBoverhangMin 1 --alignIntronMin 20 --alignIntronMax 1000000 To quantify the RNA abundance for each gene the number of uniquely mapped reads per gene was counted using the featureCounts function of the subread package (Liao et al., 2014). The read counts were normalized to transcripts per million (TPMs).

### MNase-seq analysis

A bowtie2 index comprised of the human reference genome hg19 and the sequence of the used Adenovirus strain (see section *Virus annotation* for details) was built (code available at (Schwartz and Längst, 2015)) and paired-end reads were mapped to the index using bowtie2-aligner (Langmead and Salzberg, 2012) with following parameters: --very-sensitive-local -- no-discordant. Reads were filtered to mapping quality ≥ 30 and only reads still mapped in proper pairs were kept using samtools (Li et al., 2009). Reads aligned to the Adenovirus genome were extracted for further analysis. The fragment size distribution was plotted using the CollectInsertSizeMetrics function from the Picard toolkit (https://broadinstitute.github.io/picard; Broad Institute).

### pVII positioning analysis

Fragments mapped to the Adenovirus reference with a maximum insert size of 139 bp were selected and used for pVII positioning analysis. pVII positions were called using the DANPOS2 toolkit (Chen et al., 2013). First, a pVII occupancy profile for each sample was generated and normalized to 20,000 reads using the dpos function with following parameters: -u 1e-5; -a 1; -z 5; -p 0.001, --extend 35 and --mifrsz 0. Next, the profiles were normalized by a genome wide quantile normalization to the first replicate at 0 hpi of each MNase condition (high/low) using the wiq script. Finally, pVII positions were obtained from the normalized pVII occupancy profiles using the dpos function with following parameters: -jd 50; -q 10; -z 5; -a 1.

### Rotational positioning analysis

49 to 51 bp sized fragments mapped to the Adenovirus reference of all high MNase samples were selected and pooled together. Frequencies of A/T or G/C di-nucleotides around the fragment center were calculated using the annotatePeaks.pl script from the HOMER suit (Heinz et al., 2010). Noise was filtered using the Fast Fourier Transform build in the filterFFT function of the nucleR Bioconductor package (Flores and Orozco, 2011). The cutoff was set with the option pcKeepComp=0.2. Finally, to reflect the symmetry of the pVII binding the reverse di-nucleotide frequency was added, and the total frequency was divided by 2. The same procedure was conducted to derive the di-nucleotide periodicity of nucleosomal fragments, except that 147 bp fragments of the first replicate at 0 hpi were selected.

### Ad genome accessibility

DNA fragments released under low MNase conditions were used to identify structural changes of Ad chromatin over time. Therefore, fragments mapped to the Ad genome were filtered by a maximum length of 140 bp and replicates of each timepoint were pooled. Next, all timepoints were pooled to call peaks using the DANPOS2 toolkit (Chen et al., 2013) with following parameters: dpos -m 1 --extend 35 -c 34062 -u 0 -z 20 -jd 70 -q 15 -a 1 -e 1. Fragments of each time were counted at the determined peaks using featureCounts (Liao et al., 2014) with following parameters: --fracOverlap 0.7 -p -B –largestOverlap. Peaks exhibiting in total less than 30 fragment counts were discarded. Fragment counts per peak were normalized to CPMs (counts per million). The accessibility score was obtained for each peak by fitting a linear regression on the logarithmical normalized count frequencies computed against the logarithmical infection time in min. The slope of the linear fit was used as a measure of dynamic changes of viral genome accessibility (accessibility score).

### Transcription factor motif identification

The Ad DNA sequence from position 19,890 to 19,940 was extracted and uploaded to the PROMO web server (Messeguer et al., 2002). Motif search was performed against known human and mouse motifs stored in TRANSFAC data base using the SearchSites tool. The maximum dissimilarity rate was set to 2 %. The obtained motifs were further filtered to have at least a recognition sequence of 5 nucleotides and that the corresponding transcription factor is expressed in the infected H1299 cells.

### ChIP-seq analysis

Paired-end reads were mapped against viral and human reference genomes and subsequently filtered as described in the MNase-seq analysis section. CPM-normalized coverage tracks were generated using the bamCoverage script of the deepTools package (Ramírez et al., 2016). PCA and correlation heatmap were generated using the multiBigwigSummary (50 bp bin size), plotPCA and plotCorrelation deepTools functions.

### Sites of nucleosome assembly

The R Bioconductor package nucleR (Flores and Orozco, 2011) was used to call sites of nucleosome assembly onto the Ad genome. First, fragments specific to the Ad genome were loaded into the R environment. Fragments between 137 to 157 bp from time points 1, 2 and 4 hpi of the high MNase condition were extracted for further analysis. The coverage was normalized to reads per million Ad mapped reads and noise was removed using the built-in Fast Fourier Transform (filterFFT with option pcKeepComp=0.007). Potential nucleosome positions were called based on the normalized profile using the peakDetection function with the option width=147. Peaks at genomic sites, which exhibit a mean H3.3 ChIP signal (U2OS33) of at least 2 CPM at 4 hpi were used as nucleosome assembly positions.

## ACCESSION NUMBERS

The Gene Expression Omnibus accession number for the MNase-seq, ChIP-seq and RNA-seq data reported in this paper is GSE136550. The code used to analyze the sequencing data is available at GitHub: https://github.com/uschwartz/AdVir

## SUPPLEMENTAL INFORMATION

Supplemental Information includes 8 figures, 4 movies, 2 tables.

## AUTHOR CONTRIBUTIONS

Conceptualization H.W. and G.L.; Methodology F.L., T.K., E.S., C.H., M.N., E.B., F.R., E.B., M.R., H.W. and G.L.; Formal Analysis F.L., U.S. and H.W.; Investigation T.K., C.H., F.L., C.B., E.S., M.R., H.W. and G.L., Data Curation U.S; Visualization F.L., U.S., C.B. and H.W.; Writing – Original Draft U.S. G.L. and H.W.; Writing – Review & Editing E.B., T.K., U.S., M.R., G.L. and H.W., Resources E.B., H.W and G.L; Supervision H.W and G.L; Funding Acquisition G.L. and H.W.

## ACKNOWLEDGEMENTS

Sequencing was conducted at the NGS Core of the Leibniz Institute for Immunotherapy (LIT, Regensburg, Germany) and at the Genomics Core Unit (GCU, Regensburg, Germany). The study was financed by the BMBF 01DN17003 (to G.L.), the DFG SFB960 (to G.L.), the Fondation pour la Recherche Médicale (Equipes FRM 2018 DEQ20180339229, to H.W.) and by Grants-in-aid for Scientific Research from the Ministry of Education, Culture, Sports, Science and Technology of Japan (19H04838, to T.K.). Some of the microscopy was done in the Bordeaux Imaging Center, a service unit of the CNRS□INSERM, and Bordeaux University, a member of the national infrastructure France BioImaging. HW is an INSERM fellow.

