## Supplemental data for "Adenoviral chromatin organization primes for early gene activation"

### Figure S1 Related to Figure 1

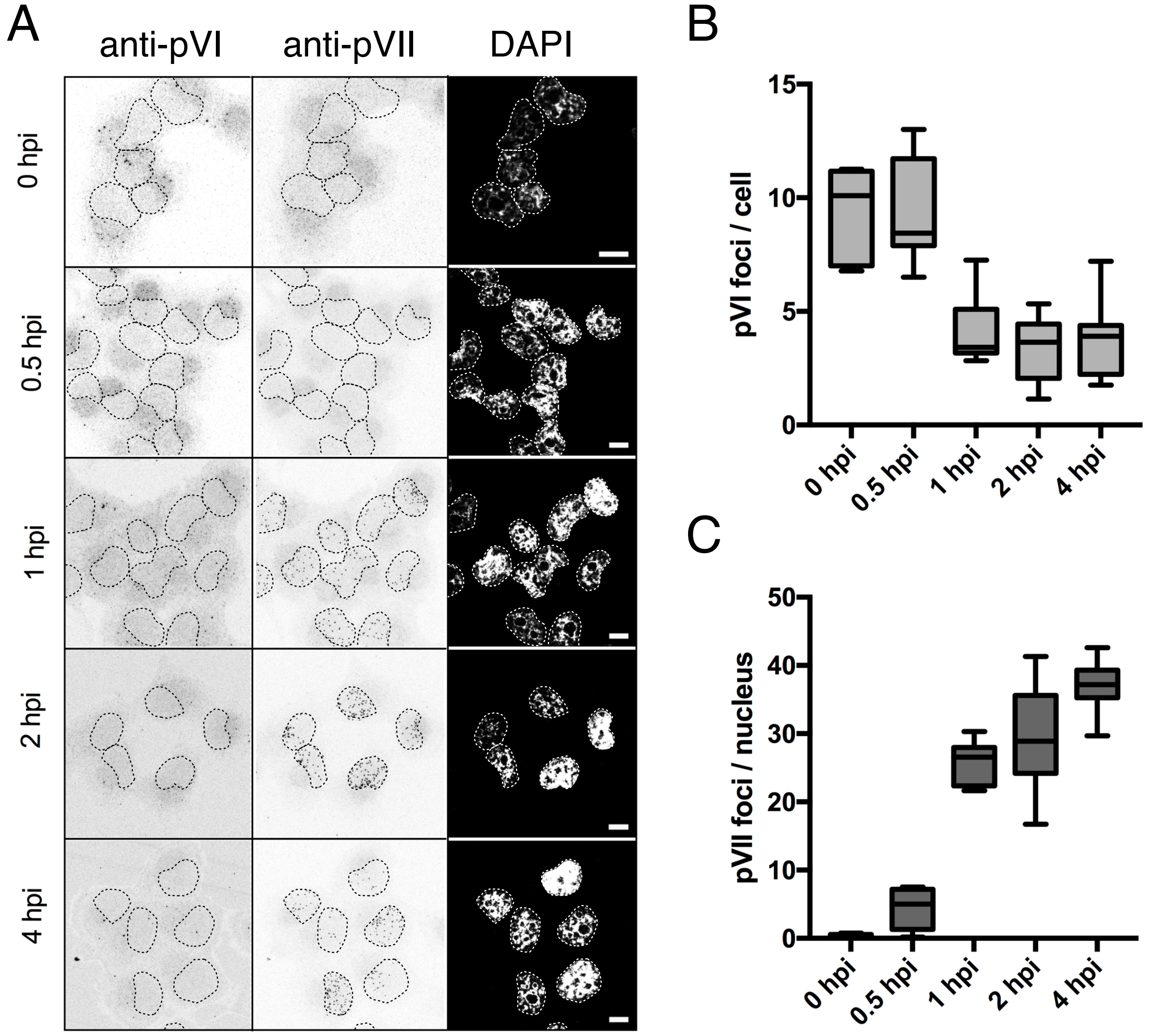


**Figure S1. Related to Figure 1**:

Spatio-temporal analysis of adenovirus infection. A) Images show representative confocal images at indicated time points stained with anti-pVI (left column) and anti-pVII (middle column) antibodies and DAPI detection of the nucleus (right column). Nuclei are indicated by dashed lines. Scale bar is 10µm. B) Quantification of the pVI signal over time as indicated on the x-axis. The distribution of pVI foci per cell are shown as Box and Whiskers plot (n > 44 cells per time point). C) As in (B) showing the quantification of pVII foci per nucleus from the same experiment (n > 44 nuclei per time point).

### Table S1 Related to Figure 1

| timepoint | replicate | sequenced reads | reads annotated to human and Ad genome | reads annotated to Ad genome | RPM annotated to Ad genome |
|---|---|---|---|---|---|
| 0 hpi | R1 | 29,121,328 | 28,484,036 | 100 | 4 |
| | R2 | 31,703,184 | 30,920,923 | 143 | 5 |
| 0.5 hpi | R1 | 23,814,664 | 22,986,503 | 32 | 1 |
| | R2 | 23,296,828 | 22,503,152 | 32 | 1 |
| 1 hpi | R1 | 24,904,732 | 24,149,420 | 50 | 2 |
| | R2 | 74,971,114 | 74,417,479 | 156 | 2 |
| 2 hpi | R1 | 27,062,105 | 25,188,583 | 382 | 15 |
| | R2 | 27,789,187 | 26,455,494 | 462 | 17 |
| 4 hpi | R1 | 18,376,378 | 17,941,436 | 45,188 | 2519 |
| | R2 | 18,670,627 | 18,303,871 | 53,942 | 2947 |

**Table S1. Related to Figure 1**:

Overview of obtained reads and alignment efficiency of RNA-seq data. RPM = reads per million mapped reads.

### Figure S2 Related to Figure 1

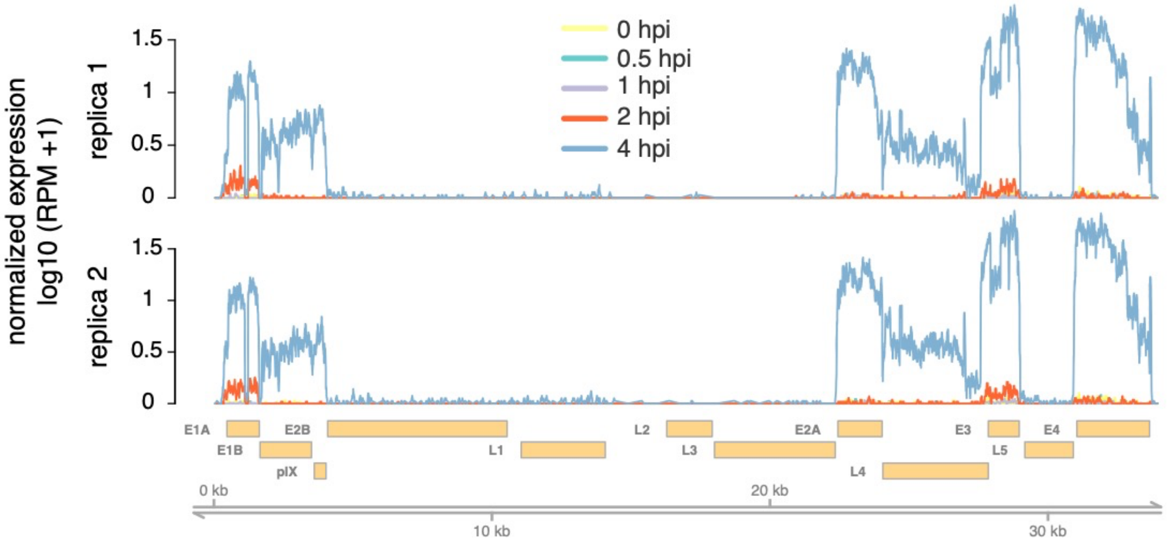


**Figure S2. Related to Figure 1**:

Profiles illustrate Ad genome coverage by RNA-seq reads. Measurements were replicated in two independent time series. Reads were normalized to RPM (reads per million mapped reads). Genomic location and underlying gene annotation of HAd-C5 (NCBI accession: AY339865.1) are indicated at the bottom.

### Figure S3 Related to Figure 3

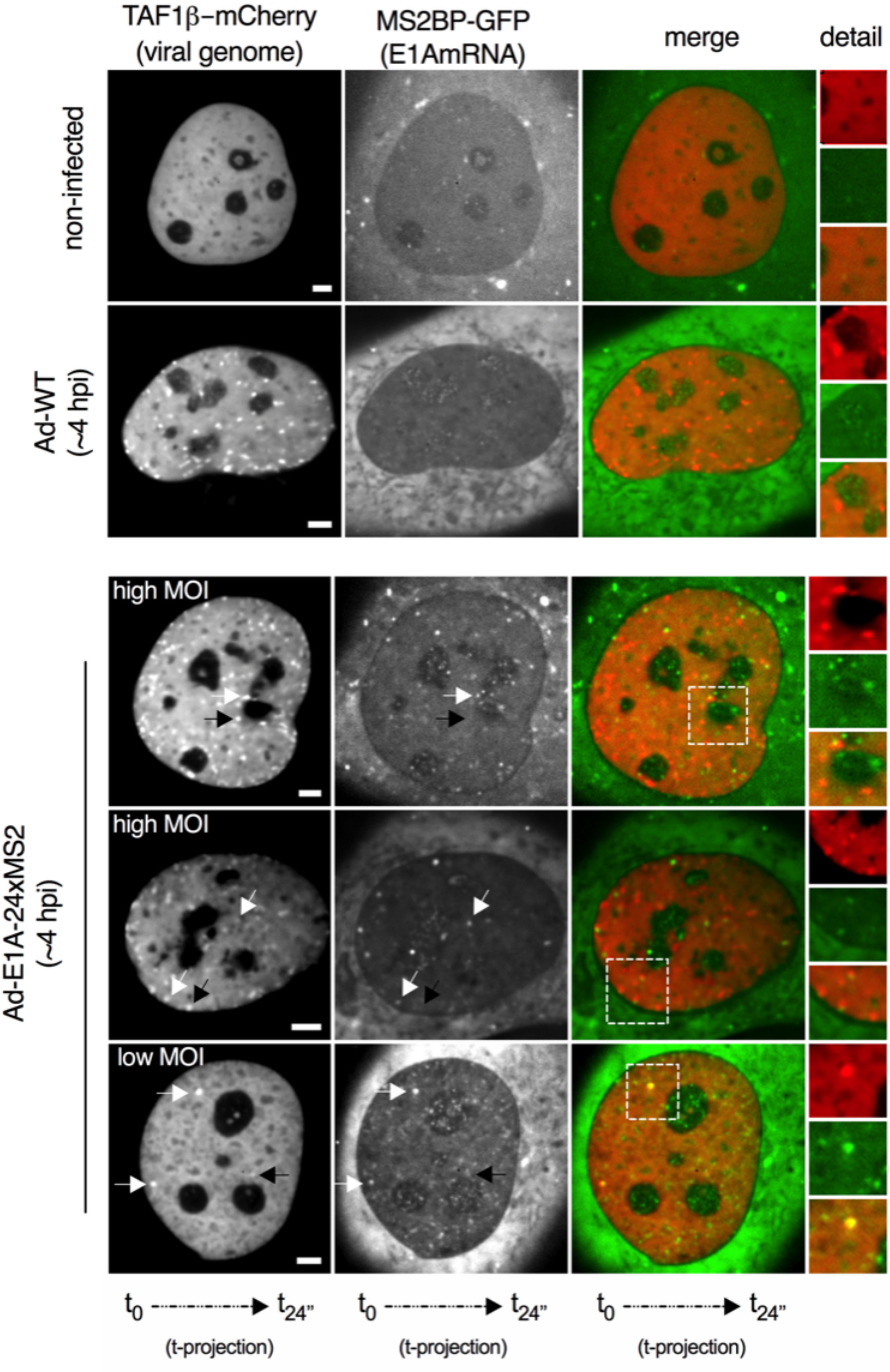

**Figure S3. Related to Figure 3**:

Representative images composed of t-projections of 24 frames imaged at ~4 hpi at a frame rate of 1 frame per second. Cells express TAF1β-mCherry showing viral genomes (first column, red signal in merge) and MS2BP-NES-GFP showing MS2-repeat tagged E1AmRNAs (second column, green signal in merge) and an overlay (third column, merge) of both channels. The first row shows non-infected control cells, the second row cells infected with untagged control virus and the third and fourth row examples of cells infected with virus expressing E1AmRNA tagged with MS2-repeats using a high MOI and the fifth row using a low MOI. Black arrows point at non-transcribing and white arrows at transcribing genomes. Details are shown as magnifications of the boxed area. Scale bar is 10$\mu$m.

### Table S2 Related to Figure 4

| MNase condition | timepoint | replicate | sequenced fragments | fragments annotated to human genome | fragments annotated to Ad genome |
|---|---|---|---|---|---|
| high | 0 hpi | R1 | 229,137,820 | 163,870,708 | 28,329 |
| | | R2 | 232,897,593 | 165,936,041 | 20,222 |
| | 0.5 hpi | R1 | 234,007,548 | 172,898,971 | 61,864 |
| | | R2 | 229,100,819 | 153,222,365 | 64,927 |
| | 1 hpi | R1 | 226,436,370 | 118,979,031 | 153,974 |
| | | R2 | 219,697,342 | 135,212,205 | 88,116 |
| | 2 hpi | R1 | 227,598,948 | 131,444,807 | 178,491 |
| | | R2 | 230,658,303 | 166,272,934 | 75,382 |
| | 4 hpi | R1 | 231,625,618 | 115,031,799 | 284,421 |
| | | R2 | 225,137,928 | 169,292,212 | 117,689 |
| low | 0 hpi | R1 | 212,388,716 | 156,852,024 | 86,603 |
| | 0.5 hpi | R1 | 218,965,404 | 153,473,717 | 205,834 |
| | | R2 | 216,033,646 | 165,531,283 | 135,786 |
| | 1 hpi | R1 | 218,399,432 | 157,676,688 | 200,931 |
| | | R2 | 216,415,263 | 142,234,881 | 541,881 |
| | 4 hpi | R1 | 212,947,907 | 178,933,555 | 445,892 |
| | | R2 | 221,396,121 | 182,403,786 | 197,983 |

**Table S2. Related to Figure 4**:

Overview of obtained read-pairs and alignment efficiency of MNase-seq data.

### Figure S4 Related to Figure 4

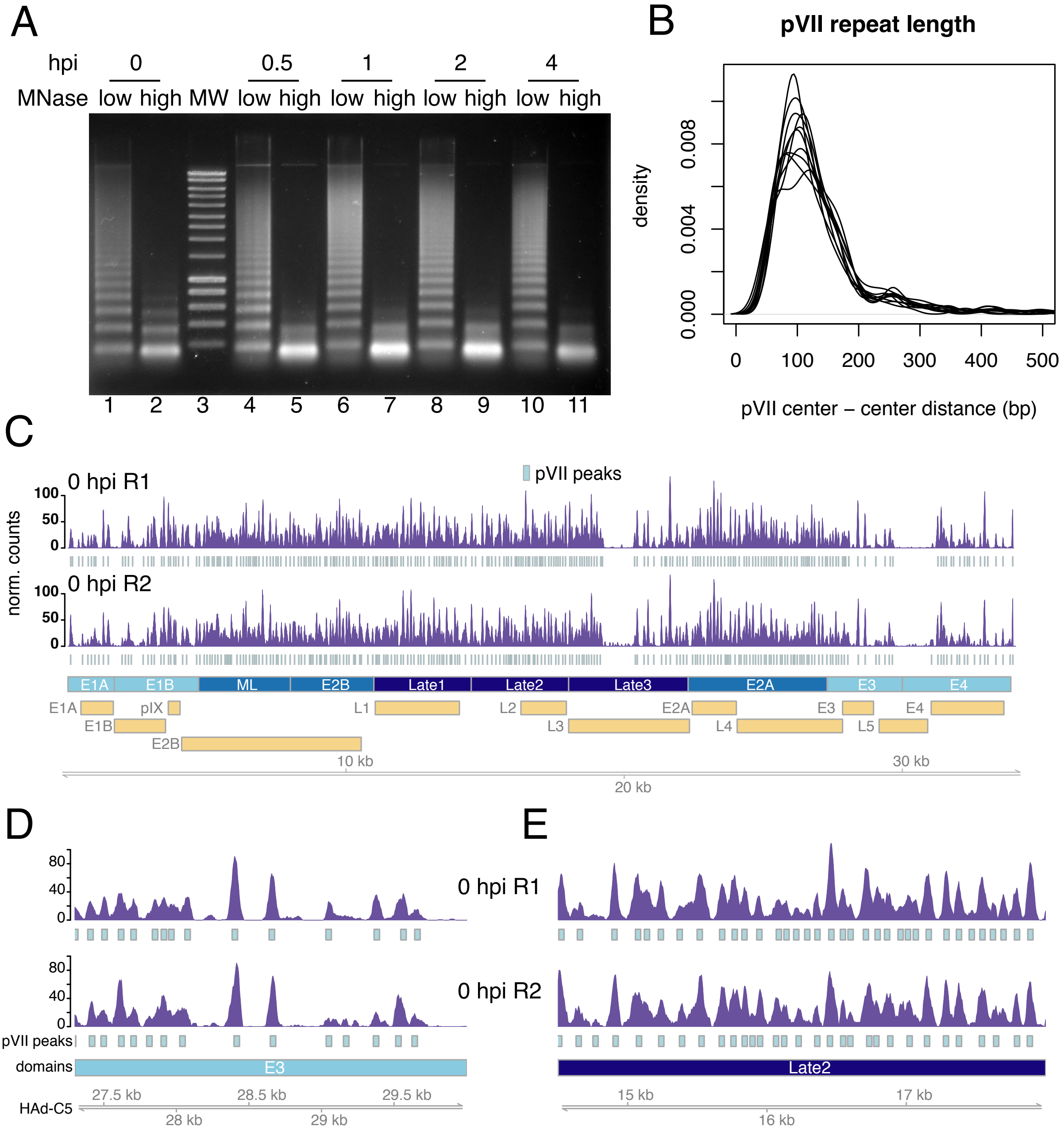

**Figure S4. Related to Figure 4**:

**A)** Nucleosomal DNA ladder after chromatin digestion of Ad infected H1299 cells using 100U (low) or 600U (high) of MNase. **B)** Distribution of pVII center-center distances of each sample along the Ad genome. **C)**, **D)** and **E)** profiles illustrate Ad genome coverage by MNase-seq reads of two biological replicates. pVII positions are represented by grey boxes. Partitioning of the genome into functional domains and the timing of their transcription is indicated by the color code (lightblue = early, darkblue=late). Snapshots of **D)** an open and **E)** a tightly packaged genomic region.

### Figure S5 Related to Figure 4

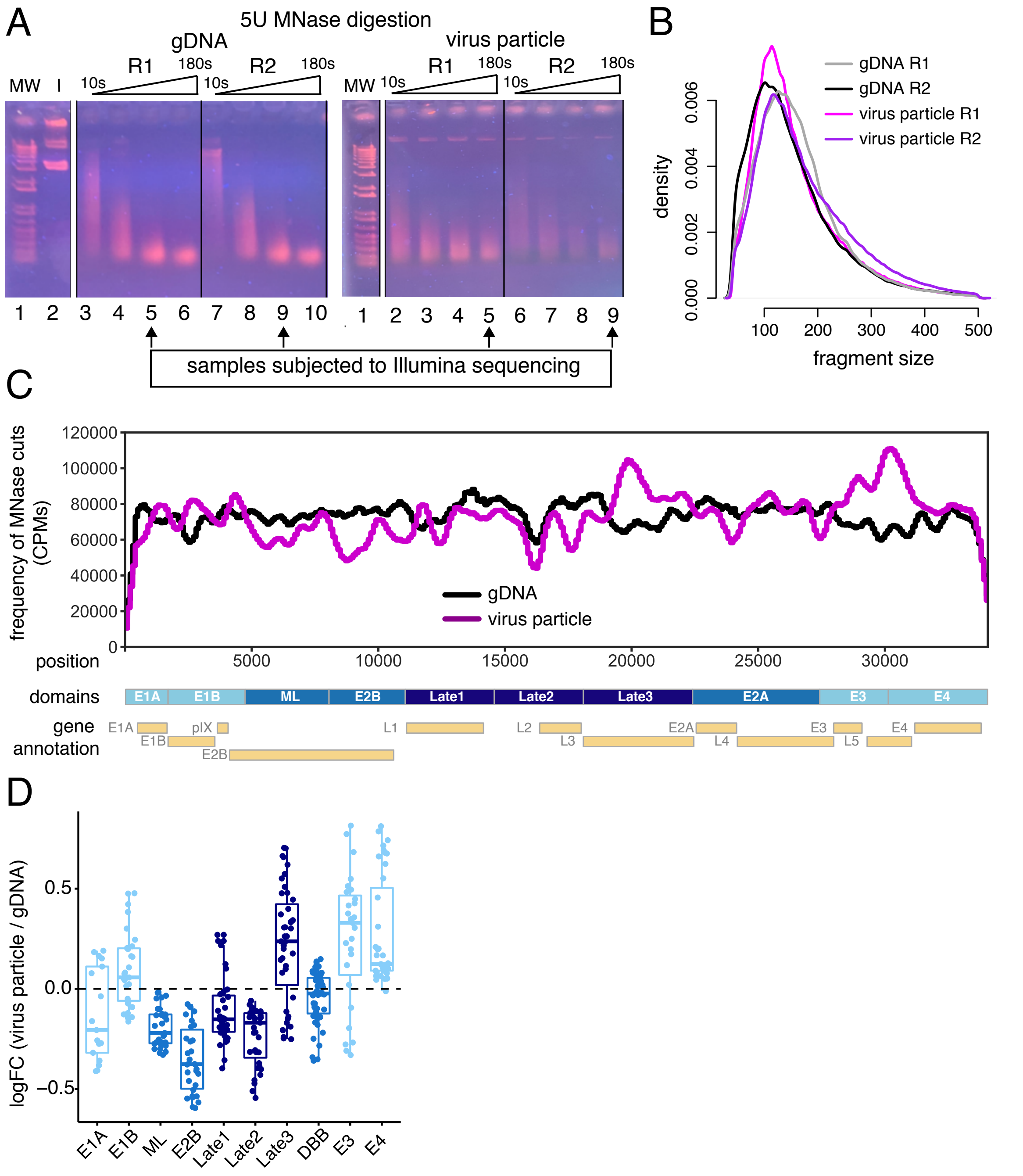


**Figure S5. Related to Figure 4**:

**A)** Agarose gel after incremental chromatin digestions (from 10s to 180s) of genomic Ad DNA (gDNA) or Ad chromatin enclosed in virus particles using 5U of MNase. gDNA was inserted into a bacterial artificial chromosome and mixed with generic DNA in ratio 1:14 (I). Virus particles were incubated for 6 min at 48°C to enable MNase infiltration. DNA from lane 5 and 9 (90s of MNase digestion for gDNA samples and 180s for virus particles, respectively) was isolated and subjected to next generation sequencing. MW = molecular weight marker; R = replicate; I = input.
**B)** Kernel density plot of the fragment size distribution of paired-end reads mapped

to the Ad genome. **C)** MNase footprinting analysis of gDNA (black) or Ad chromatin in virus particles (purple). The sites of preferential MNase cleavage were assessed as the number of 5' ends of all DNA fragments along the Ad genome. The MNase cutting frequency was normalized to counts per million sequenced fragments (CPM) and quantified in 500 bp sliding windows using a step size of 100 bp. **D)** Boxplots showing preferential regions of MNase catalyzed DNA hydrolysis in viral chromatin compared to gDNA. The data is represented as log fold change (logFC).

### Figure S6 Related to Figure 5

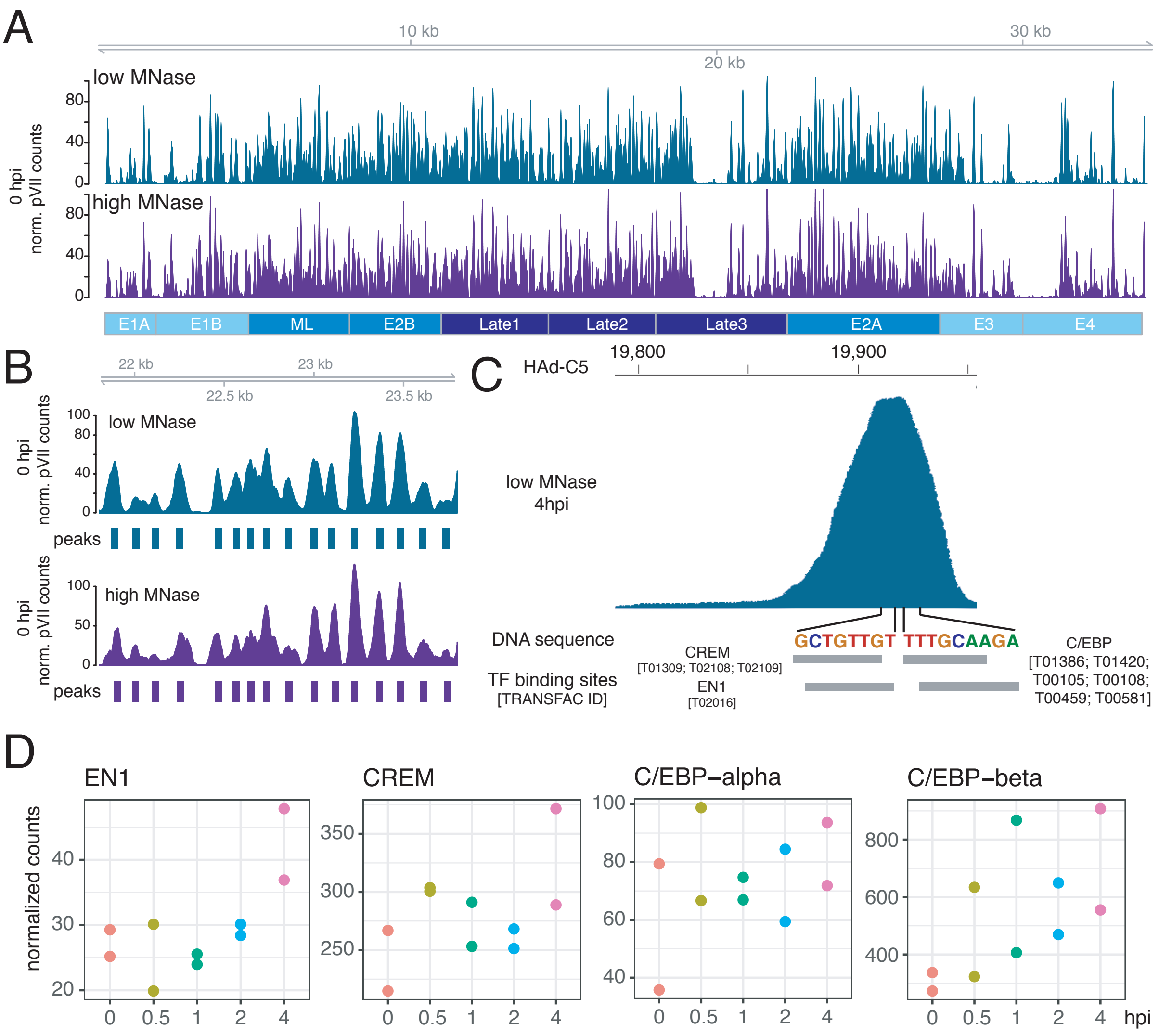


**Figure S6. Related to Figure 5**:

**A)** Comparison of pVII occupancy profile and **B)** position calling between high and low MNase digested chromatin at time point 0 hpi. **C)** Estimating the number of assembled nucleosomes onto the Ad genome. A mixed distribution of pVII and nucleosome fragments was calculated to reassemble the fragment distribution at 2 hpi and 4 hpi. The best match was obtained using 8% / 14% of nucleosomal fragments at 2 hpi / 4 hpi. **C)** Transcription factor motifs in the centre of a low MNase extracted peak at 4hpi. The genomic location is indicated at the top. The DNA sequences containing known binding motifs are highlighted. The sites of the obtained motifs are illustrated as grey box and their respective TRANSFAC ids are displayed in square brackets. D) Gene expression changes of transcription factors over the infection time course.

### Figure S7 Related to Figure 6

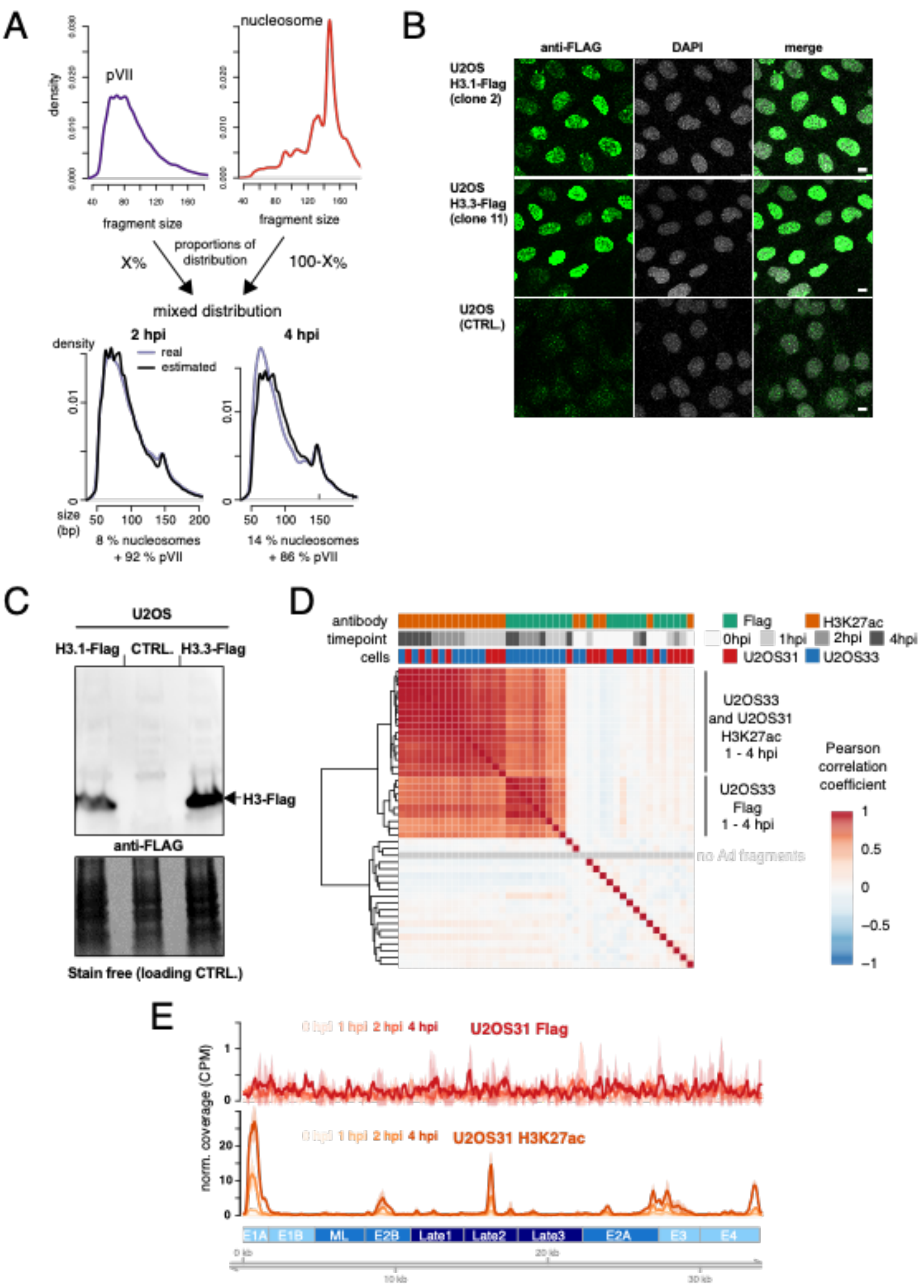

**Figure S7. Related to Figure 6**:

**A)** Estimating the number of assembled nucleosomes onto the Ad genome. A mixed distribution of pVII and nucleosome fragments was calculated to reassemble the fragment distribution at 2 hpi and 4 hpi. The best match was obtained using 8% / 14% of nucleosomal fragments at 2 hpi / 4 hpi. **B)** Images show an overview of cells from individual clones stably expressing Flag-tagged histones H3.1 (top row), H3.3. (center row) and control cells (bottom row). Cells were stained with anti-flag antibodies and detected with Alexa488 secondary antibodies and counterstained with DAPI as indicated above each column. **C)** Cells as in (B) analyzed by western blot using Flag specific antibodies. The histone H3 specific signal is indicated (arrow). A loading control is shown at the bottom. **D)** Hierarchical clustering of the Pearson correlation coefficients of ChIP-seq samples. The average fragment coverage in 50 bp bins was calculated for each sample. **E)** H3.1 and H3K27ac profiles along the Ad genome. Normalized coverage of Flag (red shades) / H3K27ac (orange shades) ChIP-seq experiments in U2OS31. The average signal of 2-3 replicates is shown and the 95% confidence interval is indicated.

### Figure S8 Related to Figure 6

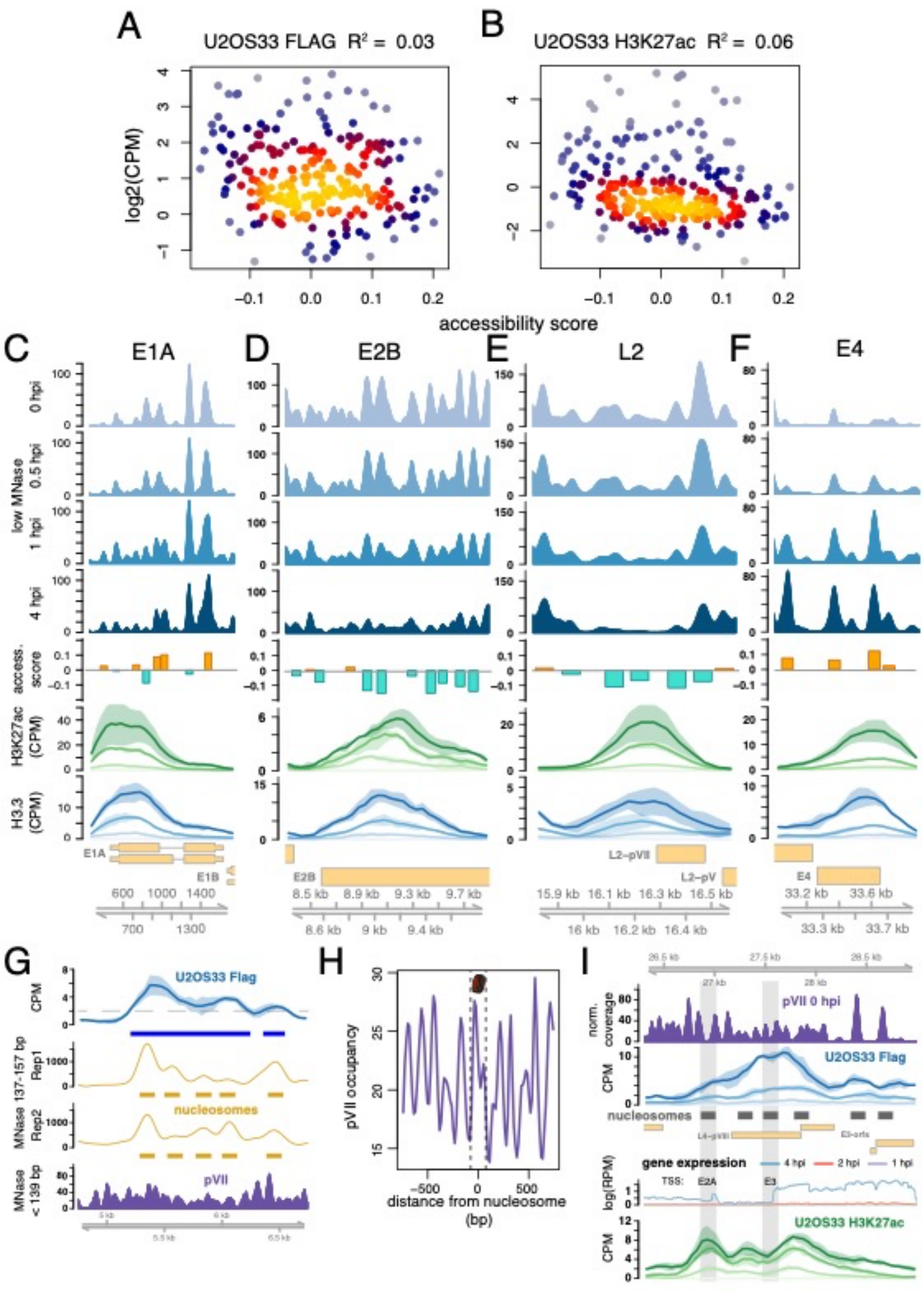

**Figure S8. Related to Figure 6**:

**A)** Scatter plots showing correlation of accessibility score with H3.3 or **B)** H3K27ac abundance at 4 hpi in U2OS33. Each dot represents one pVII peak position derived from Fig. 4D. The coefficient of determination ($R^2$) is indicated.

**C) - F)** Genome browser tracks of H3K27ac enriched sites. **G)** Genome browser tracks showing nucleosome position calling. A H3.3 coverage > 2 CPM at 4 hpi was used to define nucleosome assembly regions (blue boxes). Middle tracks depict the smoothed profile of nucleosome-sized fragments, which was used to call exact nucleosome positions (golden boxes) of both replicates at 4 hpi. pVII occupancy at 0 hpi is depicted in the bottom panel. **H)** Average pVII occupancy at 0 hpi around predicted nucleosome positions. Dashed lines indicate nucleosome boundaries. **I)** Genome browser tracks showing pVII occupancy at 0 hpi (upper panel), the H3.3 (blue shades) / H3K27ac (green shades) accumulation over time, predicted sites of nucleosome assembly (blue boxes) using nucleosome sized fragments (137-157 bp) from 4 hpi MNase-seq samples and expression levels of early genes (middle panel).
